# Mechanistic basis for understanding the dual activities of the bifunctional *Azotobacter vinelandii* mannuronan C-5 epimerase and alginate lyase AlgE7

**DOI:** 10.1101/2021.03.31.437818

**Authors:** Margrethe Gaardløs, Tonje Marita Bjerkan Heggeset, Anne Tøndervik, David Tezé, Birte Svensson, Helga Ertesvåg, Håvard Sletta, Finn Lillelund Aachmann

## Abstract

The functional properties of alginates are dictated by the monomer composition and molecular weight distribution. Mannuronan C-5 epimerases determine the former by epimerizing β-D-mannuronic acid residues (M) into α-L-guluronic acid residues (G). The molecular weight is affected by alginate lyases, which cleave alginate chains through β-elimination. The reaction mechanisms for the epimerization and cleavage are similar and some enzymes can perform both. These dualistic enzymes share high sequence identity with mannuronan C-5 epimerases without lyase activity, and the mechanism behind their activity as well as the amino acids responsible for it are still unknown. In this study, we investigate mechanistic determinants of the bifunctional epimerase and lyase activity of AlgE7 from *Azotobacter vinelandii*. Based on sequence analyses, a range of AlgE7 variants were constructed and subjected to activity assays and product characterization by NMR. Our results show that the lyase activity of AlgE7 is regulated by the type of ion present: Calcium promotes it, whereas NaCl reduces it. By using defined poly-M and poly-MG substrates, the preferred cleavage sites of AlgE7 were found to be M↓XM and G↓XM, where X can be either M or G. By studying AlgE7 mutants, R148 was identified as an important residue for the lyase activity, and the point mutant R148G resulted in an enzyme with only epimerase activity. Based on the results obtained in the present study we suggest a unified catalytic reaction mechanism for both epimerase and lyase activities where H154 functions as the catalytic base and Y149 as the catalytic acid.

**Importance:** Post-harvest valorisation and upgrading of algal constituents is a promising strategy in the development of a sustainable bioeconomy based on algal biomass. In this respect, alginate epimerases and lyases are valuable enzymes for tailoring of the functional properties of alginate, a polysaccharide extracted from brown seaweed with numerous applications in food, medicine and material industries. By providing a better understanding of the reaction mechanism and of how the two enzyme reactions can be altered by changes in reaction conditions, this study opens for further applications of bacterial epimerases and lyases in enzymatic tailoring of alginate polymers.

## Introduction

Alginate-modifying enzymes, and in particular mannuronan C-5 epimerases and lyases, can be used industrially to tailor the functional properties of alginate for food, medicine and material purposes (1–4). Post-polymerization modifications by epimerases and lyases determine the alginate applications, because they affect the viscosity and gel-forming abilities (5). Alginate is a linear polysaccharide consisting of the C-5 epimers β-D-mannuronic acid (M) and α-L-guluronic acid (G), which occur in non-random block patterns of consecutive G-residues (G-blocks), M-residues (M-blocks), and alternating MG-residues (MG-blocks) (6, 7). M-residues are present in a ^4^C_1_ chair conformation, whereas G-residues take on a ^1^C_4_ conformation. The distinct structure of G-blocks, and to a small extent MG-blocks, allows for crosslinking of chains in junction zones coordinated by divalent ions like calcium, causing gel formation (8). M/G block composition is important for viscosity and gel properties of the alginate. Mannuronan C-5 epimerases catalyse polymer-level epimerization of M-residues into G-residues (9). Alginate is harvested from brown algae, but the most studied alginate-modifying enzymes are from alginate-producing species of *Pseudomonas* and *Azotobacter* (10, 11). Two main types of bacterial epimerases are characterized: the periplasmic, calcium-independent AlgG, and the extracellular, calcium-dependent AlgEs. The AlgEs comprise seven well-studied enzymes from *A. vinelandii* (AlgE1-7) (10, 12), PsmE from *P. syringae* (13), and three recently identified *A. chroococcum* enzymes (AcAlgE1-3) (14). The AlgEs consist of from 1-2 catalytically active A-modules and 1-7 R-modules thought to modulate binding. Alginate lyases affect the viscosity of the polymer through depolymerisation. They catalyse chain cleavage through β-elimination that generates an unsaturated 4-deoxy-L-*erythro*-hex-4-enepyranosyluronate residue (Δ) at the non-reducing end (15). Different lyases show varying preferences for the four possible cleavage sites M↓M, M↓G, G↓M, or G↓G (16–18), and so lyase activity is inherently affected by epimerization patterns.

Mannuronan C-5 epimerases and lyases have been suggested to share a similar reaction mechanism (Figure 1) (15) with common initial steps of charge neutralization by residue AA1 in Figure 1, and proton abstraction from C-5 by AA2. The difference lies in the proton donation by AA3: in the epimerization reaction, it is given to C-5 from the opposite side from AA2 in the sugar ring, whereas in the lyase reaction, AA3 donates the proton to the O4 oxygen involved in the glycosidic bond and the polymer is cleaved. AlgE7, AcAlgE2, and AcAlgE3 display both activities, assumed from the same active site, highlighting the similarity between these mechanisms (16). All alginate epimerases share a common YG(F/I))DPH(D/E) motif, ^149^YGFDPHE in the AlgEs (numbering according to AlgE7) (16, 19, 20). As the bifunctional AlgEs contain the same motif, other residues must also be involved in the cleavage reaction. These determinant residue(s) could work directly as the proton donor in the lyase reaction, or indirectly by affecting a common AA3 to sometimes perform a cleavage event. The tyrosine (Y149) and the histidine (H154) in the motif are thought to be AA2 and AA3, although which role each residue performs is still not known (21, 22). For *P. aeruginosa* AlgG, which has a similar active site to the AlgEs despite low overall sequence similarity, it has been proposed that water is AA3 and histidine AA2 (22). Acidic residues around the active site could have roles in modulating the p*K*_a_-values of active residues, or they could be involved in calcium binding. E155 might stabilize the substrate carboxylate group through coordination of a calcium ion, together with D178 that is conserved in the AlgEs but not in the calcium-independent AlgG (21, 22). The calcium ion would then represent AA1. It is important to note that experiments in the absence of calcium cannot be used to probe if it actually has a catalytic role, as calcium is also required for the structural stability of the catalytically active A-modules and the carbohydrate binding module-like R-modules in the AlgEs (9, 21, 23).

**Figure 1:**
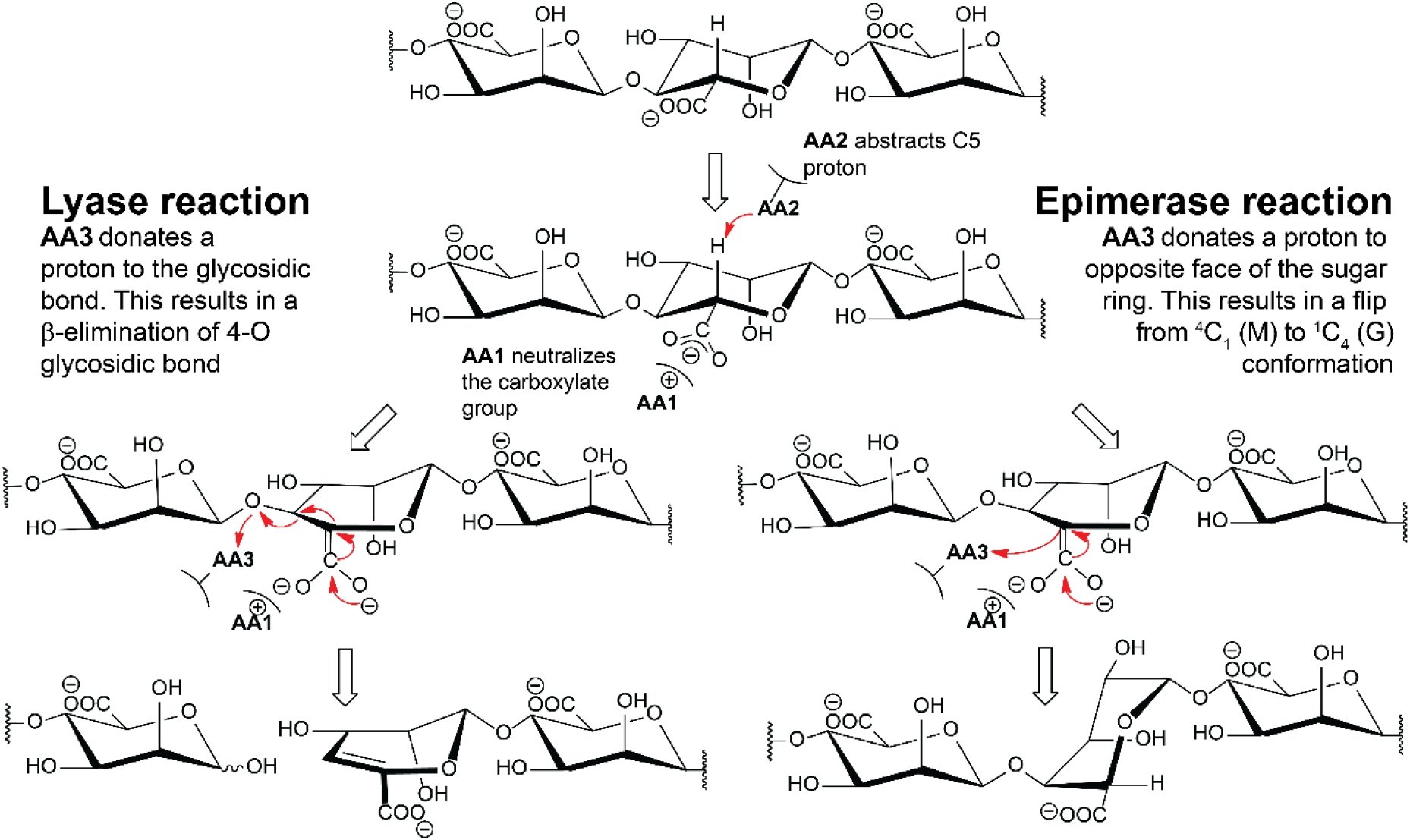
Proposed alginate lyase and epimerase reaction mechanisms. Both mechanisms start with proton abstraction from C-5 by AA2 (amino acid 2), requiring a charge neutralization of the carboxylate by AA1. In the lyase reaction, a proton is donated by AA3 to the leaving group of the glycosidic bond, whereas in the epimerase reaction a proton is donated to the sugar ring to create the epimer. Only the M-lyase reaction is shown in the figure.

There is evidence that the AlgEs form the distinct block patterns in alginate through a processive mode of action where every other sugar residue is epimerized (24, 25). Such an action would create MG-blocks from a first binding event and G-blocks from a second, explaining the occurrence of both block structures in alginate. Previous studies point to the importance of substrate interactions with charged residues far from the active site in activity and product pattern (21, 26–29).

In this study, we shed new light on the mode of action and catalytic mechanism of the bifunctional AlgE7, through investigations of the 3D-structure, substitutions of amino acids in and around the active site, and by varying the reaction conditions (pH, [Ca], [NaCl], substrates).

## Results

### The lyase and epimerization activities are mutually dependent

The epimerization pattern and preferred cleavage sites of AlgE7 have been studied previously (12, 16), and here we further elucidate the mode of action by recording ^1^H NMR-spectra at different reaction time points, and following the reaction with time-resolved ^13^C NMR. End products were verified with ^13^C HSQC (heteronuclear single quantum coherence spectroscopy) spectra (Figure 2, Figure S1, and Table S1), and HPAEC-PAD (High-Performance Anion-Exchange Chromatography with Pulsed Amperometric Detection). The investigation was performed using the two well-defined substrates polymannuronan (polyM) and polyalternating alginate (polyMG).

**Figure 2:**
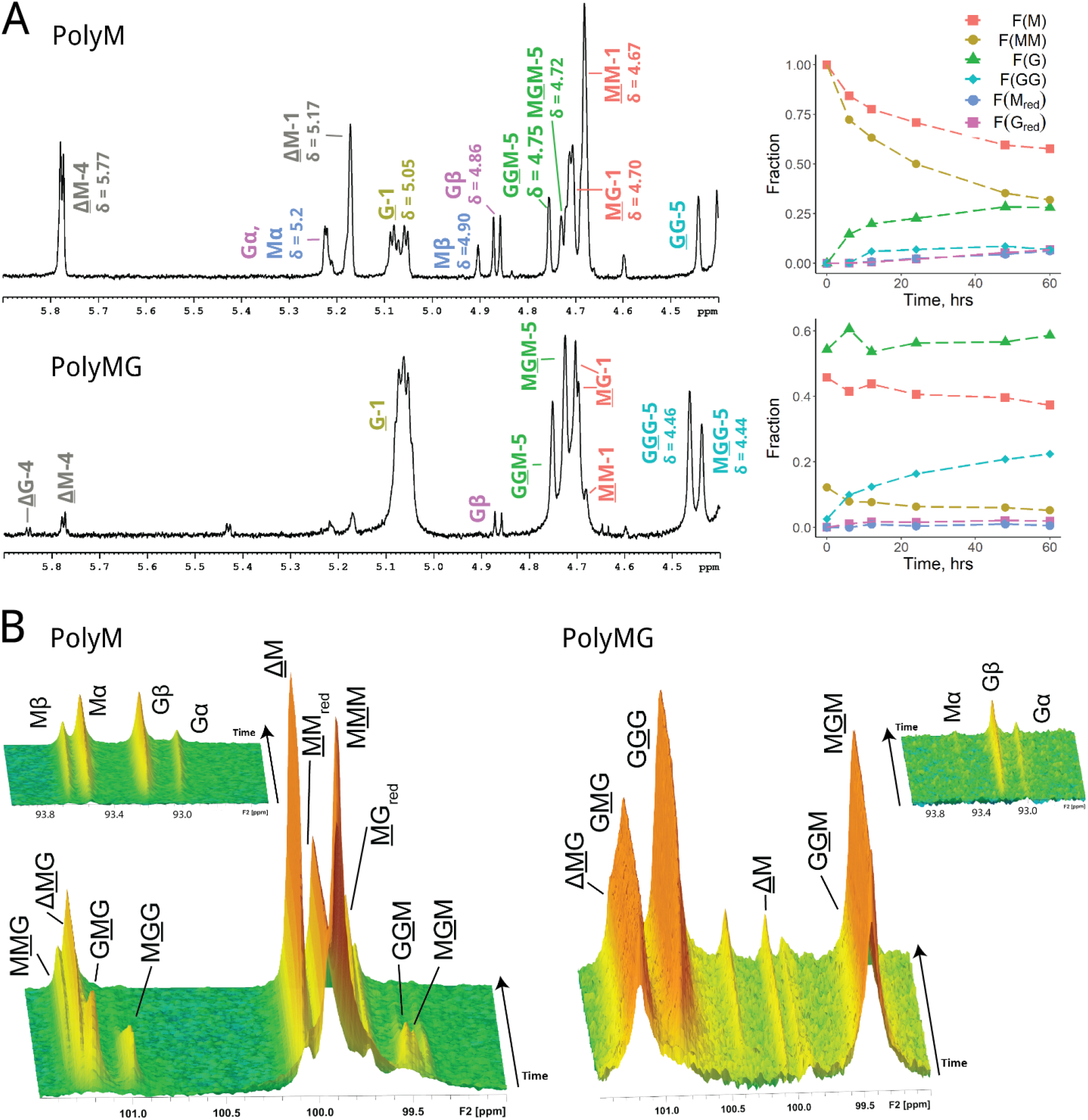
NMR spectra showing AlgE7 acting on polyM and polyMG. Underlined monomer residues give rise to the signals, and they give distinct peaks based on their nearest neighbour residues. A: ^1^H spectra after epimerization for 60 h. 5 mM HEPES, 75 mM NaCl, 2.5 mM CaCl_2_, pH 7.0. 2.5 mg/ml of substrate and 92 nM of enzyme were used in the reaction (1:300 enzyme:substrate (w/w)). Fractions of residues calculated from integration of the ^1^H-spectra at different time points are plotted to the right and shown in Table S1 (calculation of sequential parameters explained in Figure S3). B: Time resolved ^13^C-spectra of the reaction from 0-1000 min (16.7 h), recorded at 25 °C with 10 mM MOPS, 75 mM NaCl, 2.5 mM CaCl_2_ and pH 6.9. 11 mg/ml of substrate and 2.9 µM of enzyme were used (1:42 enzyme:substrate, w/w).

The ^1^H NMR spectrum of AlgE7 displayed in Figure 2A on polyM shows an absence of ΔG-signals and approximately equal amounts of M- and G-residues at reducing ends. After an initial build-up, there is a decline in G-residues, seen by the removal of M<u>G</u>G and G<u>G</u>M (Figure 2B) during the reaction course, implying that AlgE7 is able to abstract a proton from G-residues to create Δ-residues. Moreover, from the time-resolved ^13^C NMR-spectra (Figure 2B), we can infer that AlgE7 is also able to catalyse the lyase reaction (i.e. donate a proton to the 4-O) within an M-block, as it creates <u>Δ</u>M from the start. This corresponds to the potential cleavage sites G↓MM, G↓GM, M↓GM and M↓MM (the arrow denotes cleavage of the 4-O glycosidic bond). The amount of G-residues in the substrate increases during the reaction, but it is seen from the integrals of the ^1^H-spectra in Figure 2A that the formation rates of both M- and G-reducing ends remain similar for the duration of the reaction. The fact that the cleavage rates for G↓XM and M↓XM remain similar despite their relative concentrations changing drastically implies that the cleavage happens in the same binding event as preceeding epimerizations. The <u>Δ</u>M-peak is much larger than the Δ<u>M</u>G-peak, suggesting that mostly ΔMM is created, and this is supported by the HSQC-spectrum recorded at the end of the reaction (Figure S1) This is also consistent with a cleavage event happening after processive epimerizations, with a polyM block ahead. Since we see <u>G</u>G-signals in the proton spectrum after 60 h, XGGMX motifs could have become cleavage sites forming XGG-red and ΔX. No GGG signals were present in any of the spectra on polyM, indicating that there was no formation of G blocks. The absence of consecutive G-residue signals in the HSQC-spectrum at the end of the reaction supports the cleavage of G-residues.

When polyMG was used as substrate, the pattern changed. The only possible cleavage sites are G↓MG, M↓GM, G↓GM and M↓GG, as only negligible amounts of consecutive M-residues are present. The ^1^H- and time-resolved spectra show that the lyase activity decreased, only a small ΔG signal appeared, and almost no M_red_ was formed. This finding corresponds to a weak activity on the cleavage site G↓XG, which was not found on polyM, and implies a clear preference for cleaving G↓GM. From the time-resolved spectra we see that a G-block rich alginate is produced, as the MG-blocks are filled in. A total lack of activity was found on pure oligo-G, excluding cleavage inside G-blocks (Figure S2). In conclusion, the degree to which the substrate is epimerized appears to be highly correlated to the lyase activity.

The spectra shown in Figure 2A and B were recorded with 75 mM NaCl to reduce gel formation, but ^1^H NMR-spectra recorded without NaCl gave a different product pattern. This was most apparent on polyMG, where NaCl was seen to enhance epimerization and reduce lyase activity (Figure S4 and Table S1). ^13^C NMR-experiments on polyM without NaCl showed the same trends as with 75 mM NaCl, except for a total lack of MGG-signals during the reaction course (results not shown). The effect of reaction conditions was investigated in more detail and is described in a later section.

HPAEC-PAD chromatograms of the samples epimerized for 60 h in the presence of NaCl (Figure S5) showed that AlgE7 displays a pattern that is different from the M- and G-specific lyases acting on similar substrates. This reflects the more complex product patterns caused by the bifunctional enzyme. The peaks corresponding to the products created by AlgE7 overlap approximately with those of 3‒6-mers created by an M-specific lyase on polyM and polyMG, whereas the peaks do not overlap with the pure G-oligomers created by the G-specific lyase on polyG.

### Molecular basis for the lyase activity of the bifunctional enzymes and identification of R148 as a central residue

To understand the dualistic activity of AlgE7, we wanted to see how the ratio between the two activities is affected by specific amino acid substitutions. We assumed the same active site is responsible for both lyase and epimerase activities, containing the four catalytic residues Y149, H154, D152, and D178 (16, 21). Previously, a bifunctional enzyme was created consisting of the N-terminal part of AlgE7 and the C-terminal part of AlgE1 (16). The mutational study therefore focused on the N-terminal residues in the binding groove (from amino acid residue G100 to I200), as these appeared to be sufficient for lyase activity. Targeted residues are highlighted in the sequence alignment comparing the consensus sequences of the three bifunctional A-modules (ConL) and the ten epimerase A-modules from *A. vinelandii* and *A. chroococcum* (ConE) (Figure 3A, Figure S6). Charged and polar residues take part in an extensive hydrogen bonding network formed in the binding groove, illustrated with dashes in Figure 3B. Specifically, residues numbered 117, 122, 148 and 172 show differences in polar character that appear to affect positioning of the active site residues Y149 and H154 (Figure 3). Hydrogen bonding networks have earlier been shown to be important for the binding and processive action of epimerases (30).

**Figure 3:**
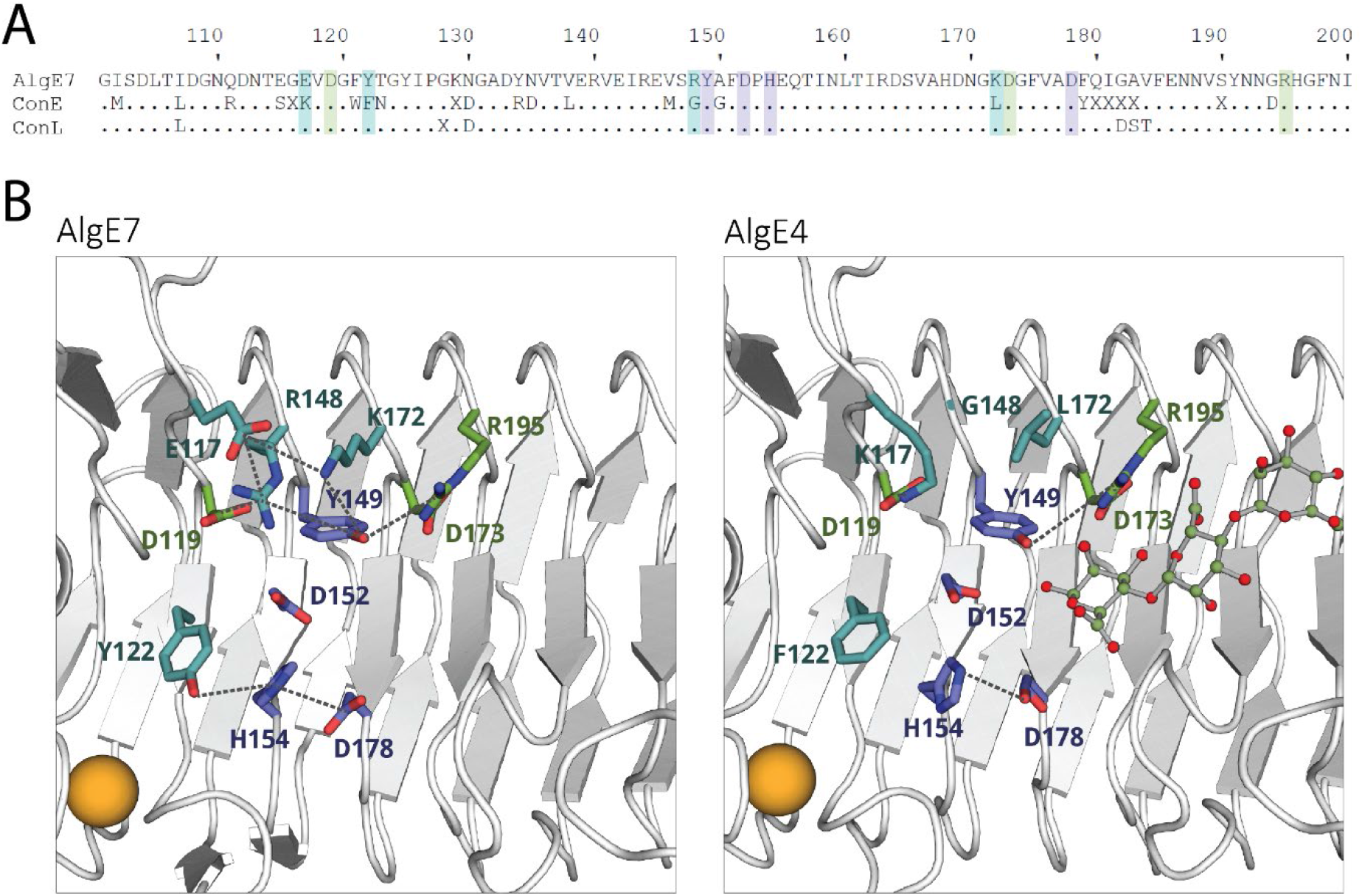
A: Alignment of AlgE7 from *A. vinelandii* with two consensus sequences. ConE is the consensus sequence of the A-modules displaying only epimerase activity from *A. vinelandii* and *A. chroococcum* (AlgE1A1, AlgE1A2, AlgE2A, AlgE3A1, AlgE3A2, AlgE4, AlgE5, AlgE6, and AcAlgE1. ConL is the consensus of the A-modules that have both lyase and epimerase activity (AlgE7, AcAlgE2A, and AcAlgE3A). Residues that are not conserved in the consensus sequence are represented in the alignment as “X”, whereas dots denote residues that are almost or completely conserved. Highlighted letters indicate amino acids replaced in this study: Purple are the catalytic amino acids, blue are the four residues differing between ConE and ConL, and green are other studied residues. AlgE7 has 82 % identity with ConL and 63 % identity with ConE. The alignment and consensus sequences are made from global protein alignments using BLOSUM 62 scoring matrix. B: The residues highlighted in A, shown in sticks in the crystal structure of AlgE4 (right) (2PYH) and a homology model of AlgE7 (left), with the same colour code as the highlights in A. Dashes are shown between residues that potentially form hydrogen bonds. The calcium ion needed for structural stability is visualised as an orange sphere. The model was built by SWISS-MODEL (48), using the structure of AlgE4 (2PYH.pdb), then energy minimized through the YASARA server (49).

Thirty-tree variants of AlgE7 with single amino acid substitutions and combinations of these were created based on sequential and visual analysis as illustrated in Figure 3. The following paragraphs will describe each group of the created variants in more detail. The activities of the variants with polyM were measured by three different methods. Firstly, release of ^3^H from [5-^3^H]-labelled mannuronan was detected in the ^3^H-assay (31, 32). This assay measures directly the first step (proton abstraction) of the reaction (Figure 1). As this step is common for both activities, the ^3^H-assay measures the total enzyme activity. Secondly, lyase activity was measured by detecting the increase in absorbance at 230 nm upon generation of the unsaturated double bonds in the resulting Δ-residues (Figure 1). The initial slope of the reaction was compared to that of the wild type. Results from the first two activity assays are shown in Table 1. Thirdly, product profiles after 48 h reaction of twenty-two of the variants were analysed with ^1^H NMR. Results from eighteen of these variants are shown in Figure 4, whereas D119A, D119N, Y122A and D173A had no detectable activity and are excluded from the figure (values are listed in Table S2). These experiments were performed using partially purified enzyme extracts where the exact enzyme concentrations cannot be determined. Therefore, the results in Table 1 and Figure 4 describe qualitative differences between the variants.

**Figure 4:**
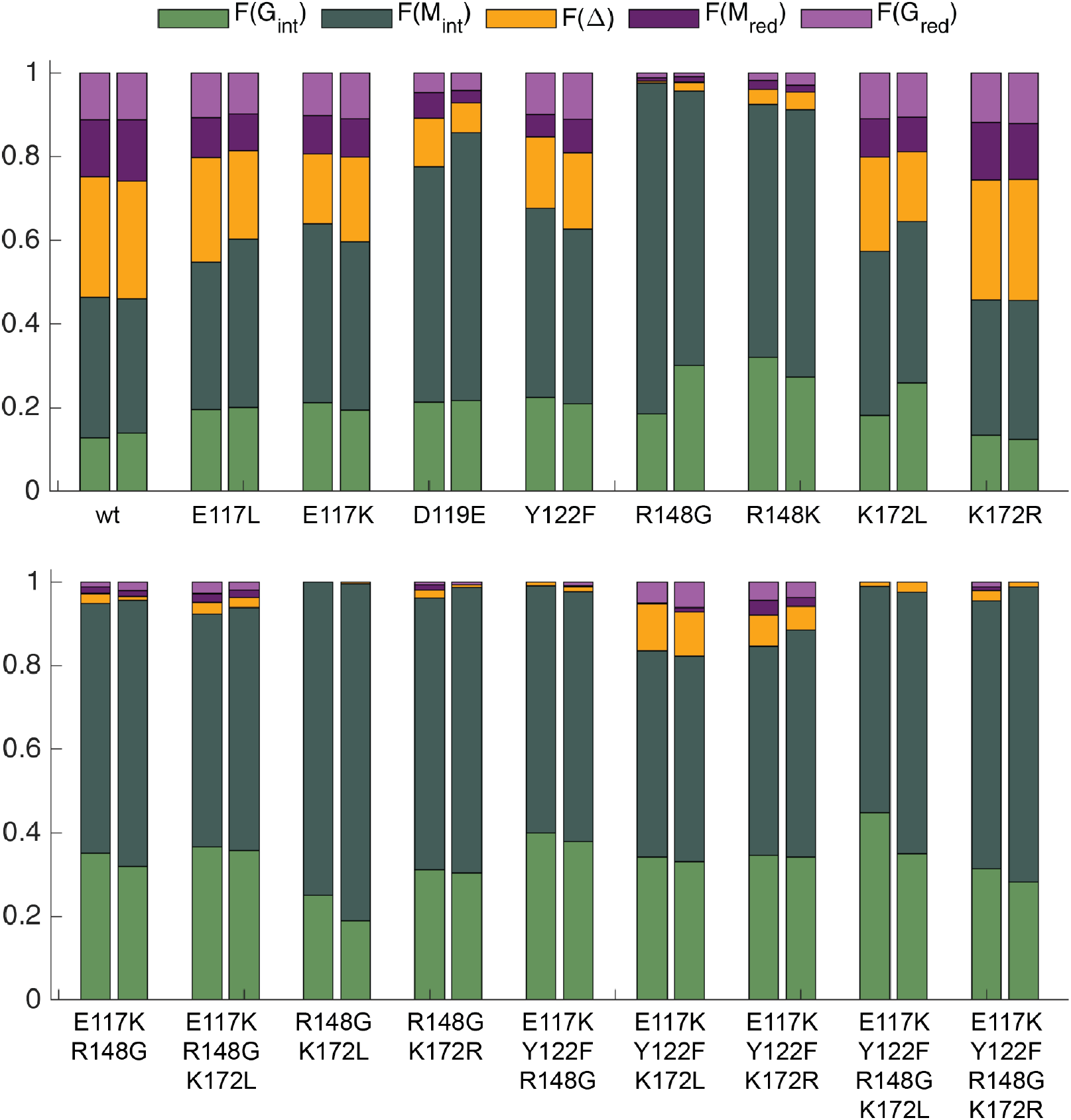
Product profiles from end point ^1^H NMR of AlgE7 wild type and mutants acting on 1 mg/ml polyM, with 2.5 mM CaCl_2_ added to the reactions, incubated at 25 °C for 24 hours. To 500 µL of reaction, 40 µL enzyme extracts were added. Each column represents one experiment, and duplicates are positioned together. Green, internal G-residues; orange, Δ-residues; dark green, internal M-residues; dark purple, M-residues at reducing ends; and light purple, G-residues at reducing ends. The sum of these fractions amounts to 1.

**Table 1:** Results from ^3^H assay and lyase assay of AlgE7 variants on polyM. All results are in percentage of wild type (wt). Bright green indicates activity between 60-100 % of wild type, dark green is activity above 100 % of wild type, light green is between 20-59 % of wild type and pale green is between 1-19 % of wild type. An asterisk (*) denotes residues involved in catalysis. ND (not determined) denotes that the experiment has not been performed. In the ^3^H assay, 0.25 mg/ml of [5-^3^H]-poly M was incubated with 2 mM CaCl_2_ and enough enzyme extract (from 5-100 µL in 0.6 ml total volume) to give comparable signals. In the lyase assay, 20 µL of enzyme extracts were added to a total of 250 µL reaction buffer containing 2.5 mM CaCl_2_ and 1 mg/ml polyM.

| Mutation | $^3\text{H}$<br>(%) | Lyase<br>(%) |
| --- | --- | --- |
| wt | 100 | 100 |
| E117K | >100 | 77 |
| E117L | ND | 92 |
| D119A | 2 | 0 |
| D119E | 9.4 | 31 |
| D119N | 0.5 | 0 |
| Y122A | ND | 0 |
| Y122F | 25 | 33 |
| R148G | 19 | 1.9 |
| R148K | 22 | 7.7 |
| Y149A * | 0.6 | 0 |
| Y149F * | 0.6 | 0 |
| Y149H * | 0.5 | ND |
| D152E * | 0.2 | 0 |
| D152N * | 2.0 | 0 |
| H154F * | 0.1 | 0 |
| H154Y * | 0.3 | 0 |
| E117K + R148G | ND | 4.0 |
| E117K + R148G + K172L | ND | 3.8 |
| R148G + K172L | 1.8 | 0 |
| R148G + K172R | 1.1 | 0 |
| E117K + Y122F + R148G | 1.3 | 0 |
| E117K + Y122F + K172L | 43 | 40 |
| E117K + Y122F + K172R | 36 | 32 |
| E117K + Y122F + R148G + K172L | 3.6 | 0 |
| E117K + Y122F + R148G + K172R | 5.6 | 0 |
| E117K + Y122F + Y149A | 1.2 | 0 |
| K172L | 76 | 77 |
| K172R | >100 | >100 |
| D173A | ND | 0.7 |
| D178E * | 0.1 | 0 |
| D178N * | 0.8 | 0 |
| D178R * | 0.3 | 0 |
| R195L | 3.7 | 0 |

Residues E117, Y122, R148, and K172 are all invariantly conserved in the bifunctional enzymes, whereas at the corresponding positions, K117, F122, G148, and L172 are conserved in the epimerases. In the epimerases, K117 forms a salt bridge with the conserved residue D119 (Figure 3). In the bifunctional enzymes, D119 is instead interacting with R148. R148 and K172 introduces additional positive charges close to the active site residue Y149. Y/F122 is the only of the four residues situated close to the catalytic histidine. We expected to suppress the lyase activity by exchanging all four residues and combinations of them into the corresponding residues found in the epimerases (E117K, Y122F, R148G, and K172L), and we wanted to see which of these residues exerted the largest effect on the activity. The variants E117L, Y122A, R148K, and K172R were created to investigate the role of the side chains further, and K172R was also included in the combination variants.

The most interesting single variant turned out to be R148G, which almost completely abolished the lyase activity while preserving the epimerase activity, both alone and in combination variants (Table 1 and Figure 4). The variant R148K retained a reduced yet significant lyase activity. Because of its significance for the lyase activity, we purified the R148G variant and performed the same ^1^H-NMR, time-resolved ^13^C-NMR and HSQC in detail characterisation as for the wild type presented above, on both polyM and polyMG (Figure 5, Figures S1-S4, Table S1). From the plot of ^1^H-NMR integrals on polyM, we see that R148G produces G-residues and GG-dyads at a slower rate than the wild type. It also forms GGG motifs that were not observed with the wild type enzyme. On polyMG it creates GGG signals only when NaCl is present (Figures 5 and S4). Also for the wild type the G-block production is increased when NaCl is present (Figure 2).

**Figure 5:**
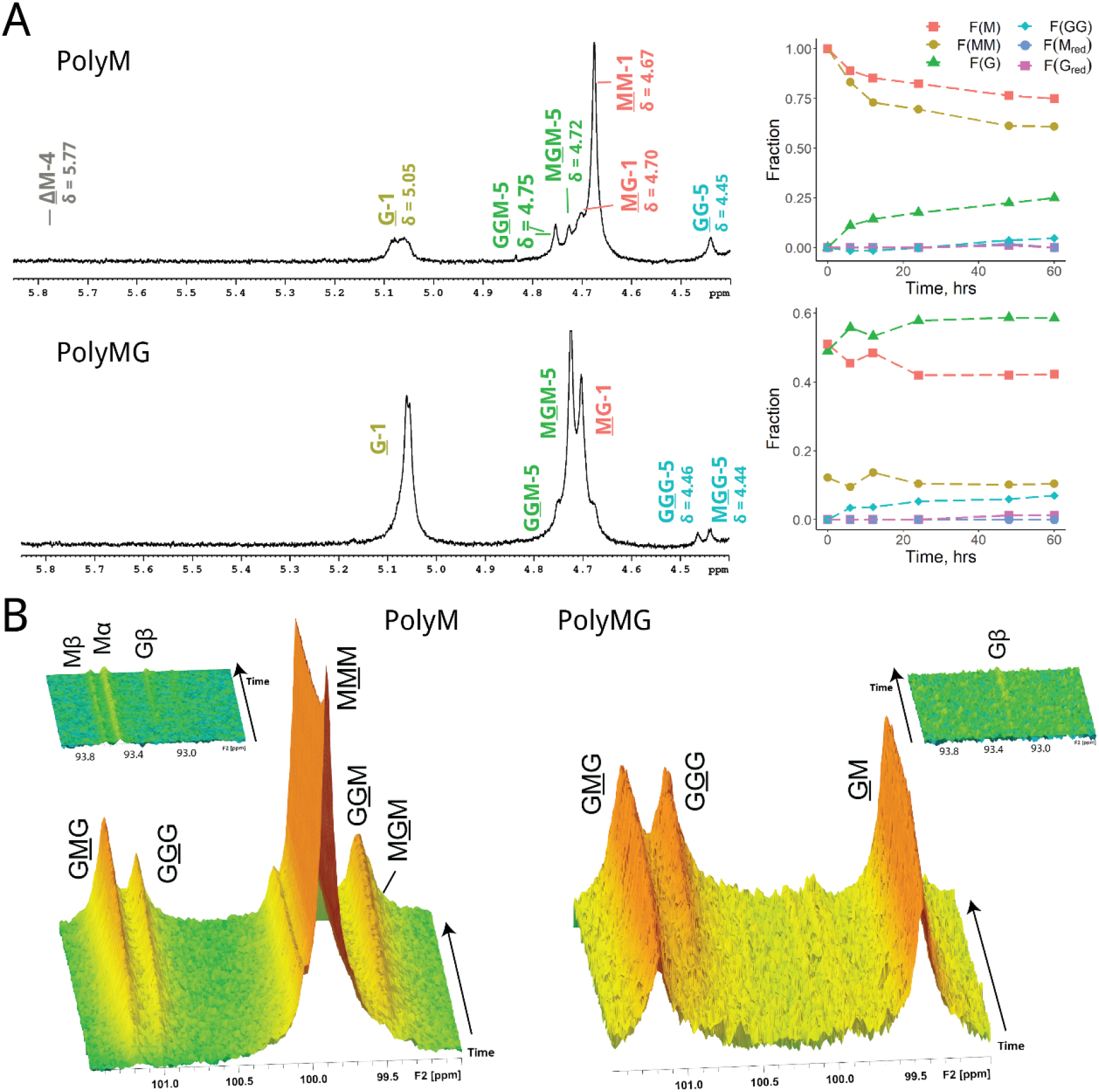
NMR-spectra showing R148G activity on polyM and polyMG. Underlined monomer residues give rise to the signals, and they give distinct peaks based on their nearest neighbour residues. A: ^1^H spectra after epimerization for 60 h, in 5 mM HEPES, 75 mM NaCl, 2.5 mM CaCl_2_, pH 7.0. 2.5 mg/ml of substrate and 0.092 µM of enzyme were added (1:300 enzyme:substrate, w/w). Plots show fractions calculated from integration of spectra recorded at different time points of the reactions. B: Time resolved ^13^C-spectra from 0-1000 min (16.7 h), recorded at 25 °C with 10 mM MOPS, 75 mM NaCl, 2.5 mM CaCl_2_ and pH 7.0. 11 mg/ml of substrate and 3.0 µM of enzyme were used (1:42 enzyme:substrate, w/w).

In contrast, E117K/L and K172L/R have lyase activity (Table 1) and product profiles (Figure 4, Table S2) similar to the wild type. Y122F has reduced activity measured by the assays in Table 1, but displays a similar product profile at reaction completion. Y122A, on the other hand, was inactive. When combining E117K, Y122F and K172L/R, the activity assays are similar to that of R148G/K, whereas the product profile shows a higher lyase- and epimerase activity. This points to indirect roles of these three residues in catalysis, as their replacements are not detrimental to the lyase activity. Nevertheless, E117L/K, Y122F and K172L appear to be connected to substrate specificity, as they produce higher amounts of G_red_ than M_red_ as opposed to the wild type which produces slightly more M_red_. The variant E117K+Y122F+K172L produces almost exclusively G_red_.

D119, D173 and R195 are conserved residues in a hydrogen bonding network around the catalytic residue Y149. The roles of these residues were investigated through the variants D119A, D119N, D119E, D173A, and R195L. D119E retained 9 % of total activity and 31 % of lyase activity, whereas D119A and D119N only retained from 0.5-2 % total activities (Table 1), implying that the charge of the side chain is more important than its size. Of the three variants, only D119E had detectable activity in the ^1^H-NMR experiment. It produced more G-residues than the wild type, which could be a result of its lower lyase activity (Figure 4). As seen in Table 1, the AlgE7 D173A variant displayed a measurable lyase activity, but too low to be detected in the ^1^H-NMR experiments. R195L had no detectable lyase activity and displayed a low total activity in the ^3^H-assay, but was not investigated with ^1^H-NMR.

To investigate the roles of the proposed catalytic acid and base in the reaction, we created the active site variants Y149A, Y149F, Y149H, H154F, and H154Y. All three variants at position 149 had a residual activity of around 0.5 % of wild type (Table 1). H154Y displayed a similar level of residual activity, whereas the activity of H154F was close to the detection limit arising from the noise. The other two proposed active site residues D152 and D178 were also investigated, through variants D152E, D152N, D178E, D178N, and D178R. D152N and D178N both display a higher residual activity than D152E and D178E, implying that for these aspartates, the size is more important than their negative charge. The D178R variant was inactive, which is not unexpected for such a dramatic change in the active site. ^1^H-NMR experiments were not performed for any of these variants.

### Complex effects on the dual activities are observed by varying reaction conditions

As already mentioned, the addition of NaCl boosted the G-block production of both wild type and R148G, especially on polyMG (Figures 5 and S4). In addition to the effect of NaCl on lyase activity, we also investigated calcium concentration, pH, and substrate concentration in more detail. The lyase activity of AlgE7 was monitored in a factorial experiment where combinations of added NaCl (0, 50, 150 and 300 mM), calcium chloride (0, 1 and 3 mM), pH (6, 7 and 8) and substrate (1 and 0.125 mg/ml) were tested. Calcium is required in the enzyme buffer to maintain structural stability of the enzymes, and a minimum of 0.16 mM calcium is present in all reactions. Lyase activity was measured with a spectrophotometric assay detecting increase in A230 upon creation of Δ-residues (shown in Figure S7), and the initial slope (Figure 6) and the end point values (Figure S8) were calculated (Table S3).

**Figure 6:**
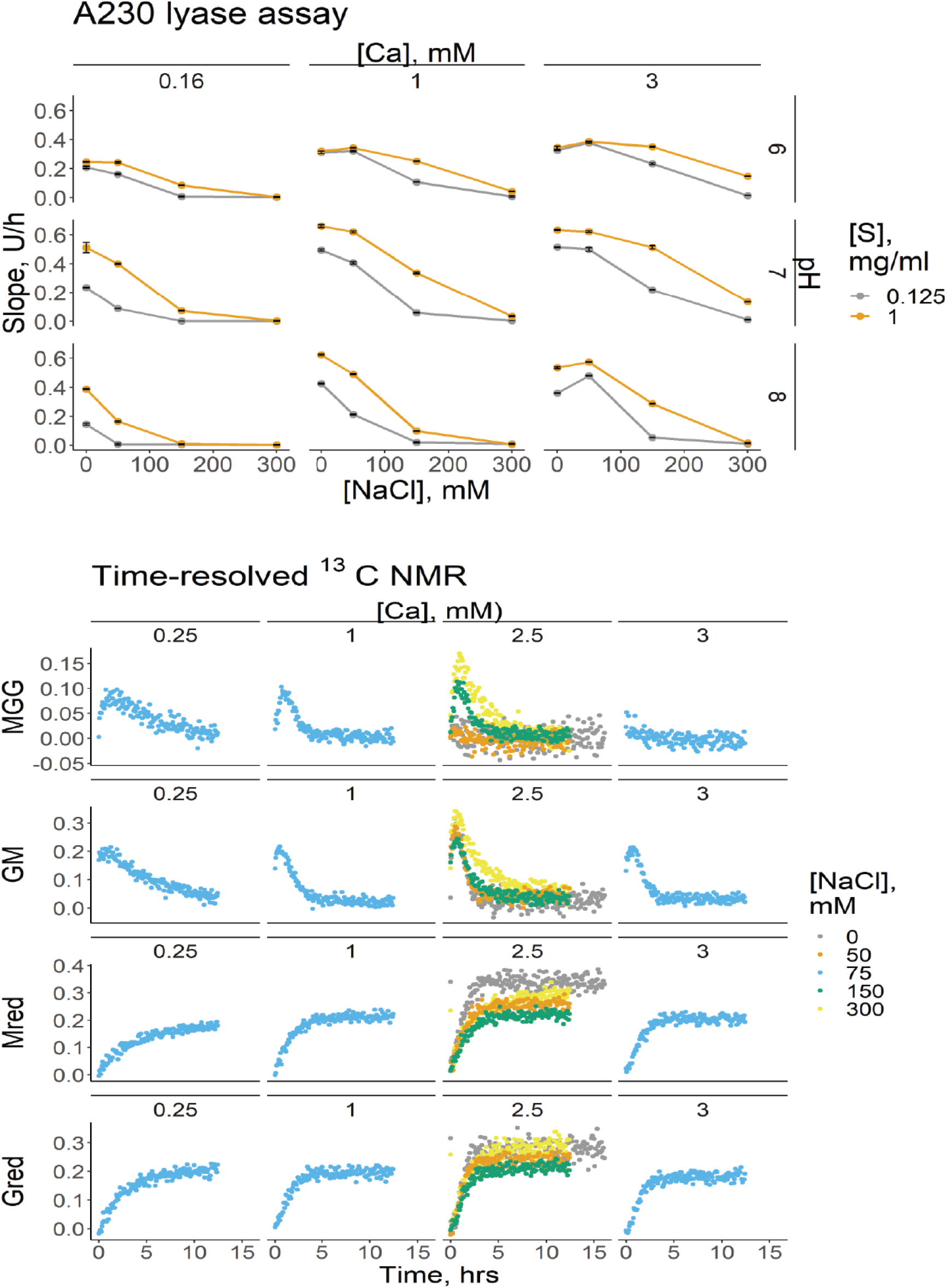
Salt dependency experiments of wild type AlgE7. Top: Slopes from a factorial spectrophotometric lyase activity assay at two different substrate concentrations [S], three different pHs, three different calcium concentrations, and four different NaCl concentrations. 0.3 µM of enzyme was added to each reaction. Slope is given in U/h, where U is absorbance units at 230 nm. Points are averages of two repetitions, and error bars show the range of values for the repetitions. Bottom: Integrated time points from ^13^C NMR with constant [S] (9 mg/ml) and constant pH (6.9) at four different calcium and salt concentrations. 3 µM of enzyme was added to each reaction. Y-axes show the relative integrals of peaks representing the given residues.

High calcium concentrations lowered the negative effect of NaCl on lyase activity, and at 3 mM calcium, the activity at 50 mM NaCl was comparable to the activity at 0 mM NaCl without supplemented calcium. The sensitivity to NaCl is also lowered with decreasing pH, although the optimal pH in general is 7. Positively charged residues are involved in the substrate interaction, as well as in the lyase reaction through R148 and K172, and pH could affect the protonation state of these residues. Activity increased with substrate concentration, except for at pH 6, 0 mM NaCl and 1-3 mM CaCl_2_, where the activity was the same at 0.125 mg/ml substrate as at 1 mg/ml. Similar trends were otherwise observed at the two substrate concentrations. The enzyme kinetics cannot be modelled by classical Michaelis-Menten kinetics, due to the continuously changing substrate from cleavage and epimerization.

To analyse the lyase and epimerase activities simultaneously, different NaCl and calcium concentrations at pH 7 with 9 mg/ml of polyM were tested with time-resolved ^13^C-NMR (Figure 6, bottom). In the experiments where the NaCl concentrations were changed, the calcium concentration was kept constant at 2.5 mM. The NMR-experiment is performed at higher substrate concentrations than the lyase assay in order to obtain a sufficient signal to noise ratio. One observed difference that could be caused by this is that the lowest lyase activity was seen at 150 mM NaCl, unlike in the lyase assay where the lowest activity was seen at 300 mM NaCl. Both the epimerase and the lyase reactions increased with calcium concentrations. However, calcium inhibited the formation of consecutive G-residues. At high calcium concentrations (2.5 and 3 mM) and NaCl concentrations below 150 mM, almost no MGG-triads were produced, unlike at higher NaCl or lower calcium concentrations where MGG-signals were produced initially and then removed through cleavage. MG and GG formation were highest at 300 mM NaCl, but this was not explained by a decrease in lyase activity.

## Discussion

### Mode of action of AlgE7 wild type

In the first experiments presented, we observed that the lyase and epimerase activities are affected by type of substrate, resulting in different product patterns. This is a challenging system for experimental investigation, as the lyase activity is affected by the epimerization, and the degradation of epimerized residues hinders the measurement of the actual epimerization activity. When *A. vinelandii* secretes the alginate used in its protective coat, the alginate has already gone through modifications by the periplasmic epimerase. AlgE7 and the other extracellular epimerases and lyases therefore encounter a substrate that is not pure polyM. The extracellular enzymes partake in forming the final alginate product in even more complex ways than what we illustrate here with AlgE7 alone, and AlgE7 itself is not expressed in large levels compared to some of the other epimerases (33). The biological role of AlgE7 has previously been indicated in release of alginate from the cell surface of *A.vinelandii* after secretion, as *algE7* knockout mutants do not release alginate to the medium (34). Newly secreted alginate contains M-residues with occasional G-residues stemming from the activity of the periplasmic AlgG, and we indeed see the highest activity of AlgE7 on polyM compared to on polyMG.

When M-rich alginate is the initial substrate, the potential cleavage sites are G↓XM and M↓XM (Figure 2), whereas on polyMG, G↓GM and G↓XG are indicated. Evidence is found that AlgE7 can cleave before both M-residues and G-residues, which gives the eight potential cleavage sites shown in Figure 7. AlgE7 has a much higher lyase activity when acting on polyM compared to polyMG, and it cannot cleave pure G-substrate. It does not create more than two consecutive G-residues on polyM before they are removed through cleavage events. To explain these findings, we hypothesize a mode of action where cleavage represents the last step of a processive epimerization, before product dissociation and release (Figure 7). Alternatively, preferred attack at the reducing end of MG-blocks flanking M-blocks and G-blocks flanking MG-blocks would give the same results. Both hypotheses explain why we see approximately equal amounts of M_red_ and G_red_ produced during the whole reaction course on polyM, even though the concentration of G-residues increases. On polyMG, on the other hand, we observe mainly G-reducing ends and almost no M-reducing ends, caused by cleavage after the formation of G-blocks. The hypotheses also explain why we observe <u>Δ</u>G-signals only from the MG-substrate, because assuming AlgE7 moves relatively to its substrate in the same direction as AlgE4 (30), it could only create those signals on polyM in the unlikely event that it reaches the end of another MG-block. More <u>Δ</u>MM is produced than Δ<u>M</u>G from polyM (Figure 2B), and on M-rich alginate, attacking the reducing end of G- and MG-blocks would likely produce ΔMM. The fact that the enzyme cannot cleave pure G-alginate also fits into this hypothesis, as this alginate cannot be epimerized. The exact positioning of various substrates might determine if epimerization or cleavage happens, which could be determined by substrate features. MM-residues are more outstretched in the chain than alternating MG or GG-residues, and span around 10.5 Å, compared to around 8.7 Å for GG-residues (35). In addition, M residues have both a hydrophilic and a hydrophobic face, whereas G residues lack a hydrophobic face. They also have a different configuration of all the polar groups around the ring carbons, and due to their distinct chair conformations (^4^C_1_ for M and ^1^C_4_ for G), they result in different three-dimensional structures of the alginate chain.

**Figure 7:**
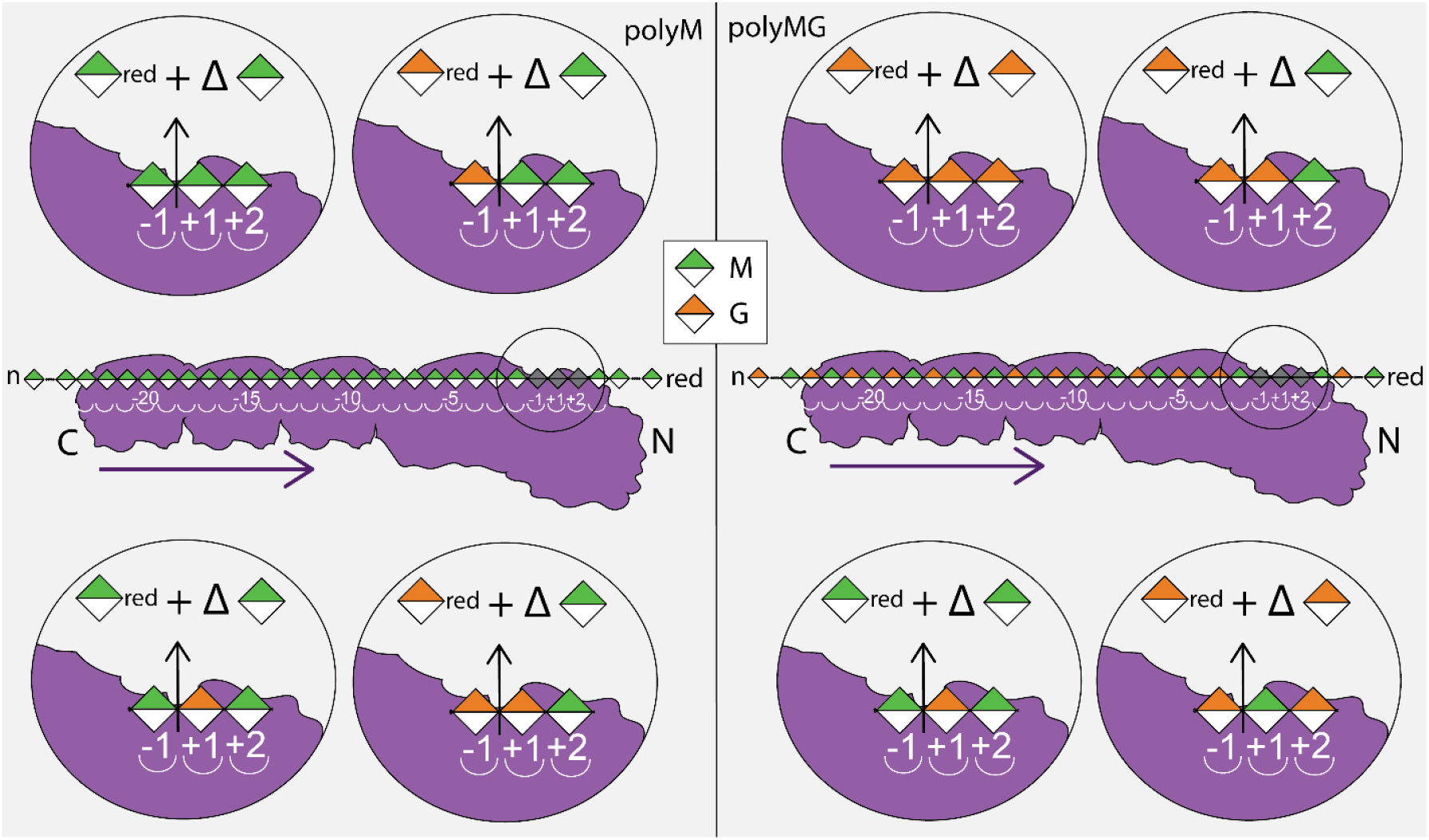
Illustration of the possible cleavage sites from the observed product patterns of AlgE7 on polyM (left) and polyMG (right). In the middle figures, AlgE7 is shown as it binds to the substrate it is presented with initially, but this will change during the reaction course. Except for M↓MM, all cleavage sites are in accordance with cleavage after a processive binding event, or due to preferred attack of the end of MG- and GG-sequences. For polyM, we observed creation of M_red_, G_red_ and ΔM, which can be caused by cleavage at the four cleavage sites illustrated in the left figures. On polyMG, mainly G_red_ and ΔM are observed, corresponding to cleavage site G↓GM. However, we also observe small amounts of M_red_ and ΔG, and four cleavage sites shown in the right figures are possible on polyMG. Sugar residues are coloured according to the Symbol Nomenclature for Graphical Representations of Glycans (50). The purple arrows indicate the proposed direction of movement for the processive AlgE4 (30). AlgE7 consists of one A-module and three R-modules, and the outline of all four modules is shown. The spatial arrangement of these modules is suggested in the figure to be elongated, but this is not studied experimentally.

### The enzymatic mechanism of epimerization and cleavage

The substrate might affect whether epimerization or cleavage happens, but the reason why only some of the AlgEs are bifunctional is due to sequence determinants in the enzymes. As three of the four important determinants (E117, R148, and K172) are positioned on the side of the active site tyrosine Y149, we propose that H154 is the proton abstractor common in both reactions (AA2 in Figure 1) and Y149 is the catalytic acid (AA3) (Figure 8). The orientation of a ligand observed in the crystal structure of AlgE4 is with the reducing end towards the N-terminal of the protein, and this has also been deemed as the most likely orientation during processive action through molecular dynamics simulations (21, 30). This orientation seemingly rules out that Y149 could abstract the H-5 protons from M residues, illustrated in Figure 8. In the figure, a sugar residue is modelled in the active site based on an elongation of the crystal structure ligand, placing the H-5 in an angle that is accessible for H154 but not for Y149. After abstraction of H-5, on the other hand, the opposite plane of the sugar is available for proton donation by Y149. The exact positioning of Y149 relative to the substrate could govern its donation either to C-5 or O-4 of the residue in subsite +1. This illustrates how epimerization at C-5 gives rise to significant differences in how the ligand is placed in the binding site. We can therefore not be sure if the same residue performs proton abstraction of both M-residues and G-residues, as the positioning would be different. Alginate lyases that can cleave G-residues can perform an *anti* β-elimination using either a histidine or a tyrosine as a base, and a tyrosine as an acid (36, 37). Based on this, it cannot be ruled out that Y149 might perform the role of the base when the enzyme removes G-residues, as it might be in a better position for this than the histidine (seen in the last step of epimerization in Figure 8). The two active site aspartates D152 and D178 have been proposed to be associated with substrate stabilization in AlgE4 (21), and D178 could also neutralize the charge of the carboxylate group through the coordination of a calcium ion (22). For simplicity we have not shown the neutralization of the carboxylate charge in Figure 8.

**Figure 8:**
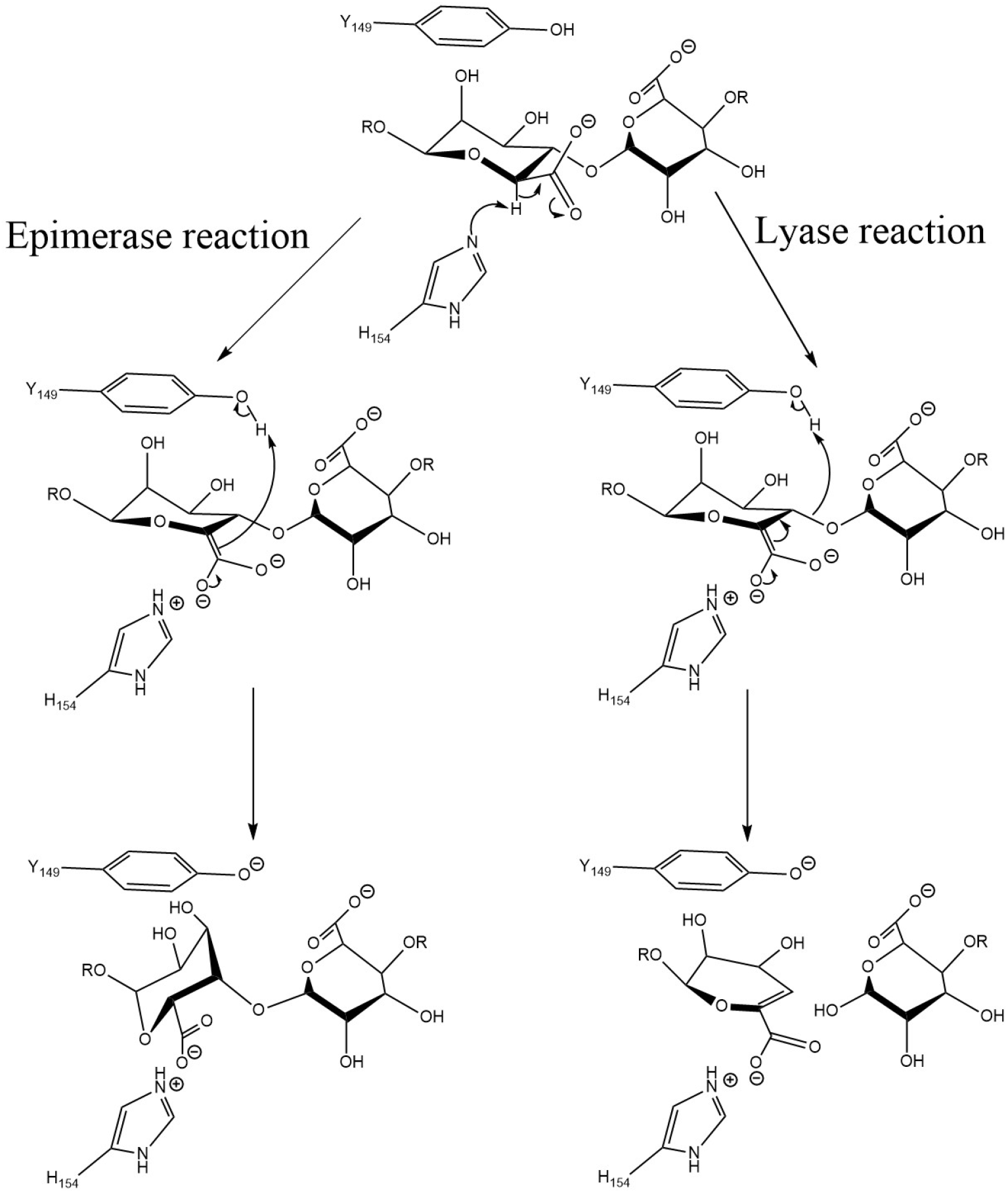
Proposed mechanisms of epimerization and lyase reaction of AlgE7 on M-residues (XMMX substrates).

R148 was shown to be the most important residue for the lyase activity of AlgE7, as the lyase activity almost completely disappeared in the R148G variant. Based on the result that the R148K variant retains slightly more lyase activity than R148G, the charge of R148 seems to be important, whereas the guanidino group of the arginine is required for optimal activity (Table 1 and Figure 4). We have considered the possibility that R148 donates the proton to the glycosidic bond itself, but it is located too far from the glycosidic bond in the proposed reaction mechanism (Figures 3 and 8). We propose that R148 is essential in influencing whether Y149 donates the proton to the other face of the sugar ring (epimerization), or to the glycosidic bond (lyase reaction), and that without it, only the epimerization happens.

The role of E117, R148 and K172 might be to position R148, Y149 and the substrate residue in the active site through electrostatic interactions, as well as perturbing the p*K*_a_ of groups involved in catalysis. Even when combining E117K, Y122F, R148G, and K172L substitutions, a minute lyase activity can be observed in the NMR-spectra (Figure 4), suggesting that these residues are involved in the lyase reaction only indirectly. The effects of the variants E117K, Y122F and K172L on the lyase activity was not as large as for R148. They did, however, produce more G-residues than M-residues at reducing ends, unlike the wild type that produced similar amounts. The variants therefore have a preference for G↓MM over M↓XM. This result highlights the sensitive fine-tuning of interacting residues relative to the positioning of the specific substrate in catalysis. E117 and K172 are both situated a bit away from the putative substrate binding site, and likely affect substrate binding indirectly through the hydrogen bonding network that includes R148 and Y149 (Figure 3). As Y122A was inactive, the aromaticity and/or the size of this residue is essential. Similar results were obtained for AlgE4, where the F122Y variant retained 65 % of the wild type activity whereas the F122V variant only retained 0.2 % (21). AlgG has a tyrosine positioned in the corresponding position, and although the alanine variant of this residue was not detrimental, it reduced the activity to 20 % of the wild type activity compared to 66 % for the phenylalanine variant (22). For AlgG, it was concluded that this residue has a role in coordinating the active site histidine. We also propose that it has a role in correct positioning of the substrate for lyase activity.

The hydrogen bonding network between the charged residues around the active site is important not only for the catalytic mechanism, but also for the processive mode of action of the AlgE epimerases. D119A, D173 and R195 are conserved in the block-forming AlgE epimerases but have no equivalents in AlgG that introduces only single G-residues to alginate chains. Substitutions of these residues that remove the charges are detrimental or at least diminishes activity dramatically, as seen here for AlgE7 (Table 1) and previously in AlgE4 (21, 30). Block-formation requires either a processive action with the ability to stay attached to the substrate for consecutive catalytic events, or at least a preferred attack with the ability to bind specifically to already epimerized alginate chains. The strength of the substrate interaction would be essential in both scenarios and could be the property that is affected when D119, D173 and R195 are mutated.

High NaCl-concentrations inhibited the initial lyase activity in the spectrophotometric lyase assay but not in the time-resolved NMR-experiments (Figure 6), which could be explained by the higher substrate concentrations of the latter (9 mg/ml, against 0.125-1 mg/ml in the lyase assay). Na^+^ and Cl^-^ ions shield charged residues in the substrate and enzyme that are essential in substrate interaction. A high substrate concentration would reduce the requirement for a strong enzyme-substrate interaction, as new substrate chains would be easily available for binding. Charged residues (E117, R148 and K172) are also implicated in the catalytic reaction, but if shielding of these were detrimental, we would not expect that a high substrate concentration would diminish the effect from NaCl. The loss in G-block formation at high calcium concentrations, but only at low salt concentrations, probably stem from sodium ions shielding the charges in the MG-substrate, preventing it from forming junction zones with calcium that would render it inaccessible to subsequent G-block formation. We see that NaCl promotes formation of G-blocks for both wild type and R148G (Figure 2, Figure 5, Figure S4), and as the same result is seen for the lyase-inactive variant, it must be caused by an increase in epimerization activity and not only a change in lyase specificity. At the low substrate concentrations in the lyase assay, this effect would be smaller, and here calcium is instead seen to save the activity at high NaCl concentration. It would be interesting to further investigate the effects from different substrate concentrations, in combination with an investigation of residue or module determinants of the substrate interaction and processive mode of action.

## Conclusions

In summary, the results presented here demonstrate what was previously suggested, namely that the epimerisation and lyase activities in the bifunctional AlgE7 share the same catalytic site. The two activities affect each other by constantly changing the alginate chains, and the product formation is highly dependent on which substrate the enzyme is acting on. The lyase preferential cleavage sites are found to be M↓XM and G↓XM when using the well-defined alginate substrates polyM and polyMG. The mechanistic insights obtained through amino acid replacements and structural analysis suggest a likely catalytic reaction mechanism on M-residues, where H154 acts as a catalytic base (AA2) in the reaction, and Y149 takes the role as a catalytic acid (AA3). The variant R148G, positioned next to the catalytic residue Y149, abolished almost all lyase activity but retained epimerisation activity, and is thought to affect whether Y149 donates the proton to the glycosidic bond or to the other side of the sugar ring. Cleavage of both M- and G-residues is evident from the identified reaction products. The reaction conditions affect the enzymatic activity, where higher NaCl reduces the lyase active and increase G-block formation, whereas high calcium concentrations increase lyase activity and decrease G-block formation. The findings in this study explain the mechanistic basis behind the dual activity of AlgE7, in comparison to the AlgEs with mainly epimerase activity. New insight into how the two activities can be modulated relative to each other, both in terms of amino acid residues and reaction conditions, will aid in future applications of the AlgEs on tailoring of alginates.

## Materials and methods

### Alginate substrates

All alginate substrates were freeze dried and weighed out upon use. Polymannuronan (polyM) was produced from the epimerase deficient AlgG *Pseudomonas fluorescens* strain PF20118 (NCIMB 10525) (38). This substrate has an F(MM) of 1.00, and a degree of polymerization (DP_n_) of around 370 determined from its molecular weight averages with Size-exclusion chromatography with multi-angle light scattering (SEC-MALS) (M_n_ of 73 kDa and M_w_ of 146 kDa) (Aarstad et al. 2019). ^13^C-1 labelled polyM was produced from fermentation of the same strain using ^13^C-1 (99 % ^13^C) D-fructose (Cambridge Isotope Laboratories, USA). This substrate was further subjected to acid hydrolysis as described elsewhere (39, 40), and had a final DP_n_ of around 70. [5-^3^H]-labelled mannuronan was produced by growing the same strain on [5-^3^H]-labelled glucose (Amersham). PolyMG was produced by epimerizing polyM with AlgE4 epimerase (41), until reaction completion, obtaining a final F(G) of 0.46 and F(GG) of 0. ^13^C-labelled polyMG was created in the same way, using ^13^C-polyM. This substrate was subjected to acid hydrolysis, obtaining a DP_n_ of around 80. ^13^C-labelled OligoG was produced by *in vitro* epimerization of ^13^C-labelled polyM with AlgE1 epimerase, and then subjecting this product to acid hydrolysis (4, 42) to obtain a final DP_n_ of around 21 and an F(G) of 0.97.

### Mutagenesis, protein production and purification

Plasmids encoding AlgE7 variants (Table 2) were constructed based on wild type AlgE7 (plasmid pBG27) (12) using the QuikChange™ Site-directed Mutagenesis Kit (Stratagene/Agilent) or Q5 site directed mutagenesis (New England Biolabs), according to instructions by the manufacturers. Primer sequences can be provided upon request. All cloning was performed using *E. coli* DH5α competent cells (New England Biolabs).

**Table 2:** Plasmids used in this study. Sequences can be provided upon request.

| Plasmids | Description | Reference |
| --- | --- | --- |
| pTYB1 | Expression vector | New England Biolabs |
| pBG27 | pTRc99a derivative, contains *AlgE7* gene | (12) |
| pKK2 | pBG27 with *AlgE7*-E117L | This study |
| pRS16 | pBG27 with *AlgE7*-E117K | This study |
| pRS38 | pBG27 with *AlgE7*-D119A | This study |
| pRS39 | pBG27 with *AlgE7*-D119E | This study |
| pRS40 | pBG27 with *AlgE7*-D119N | This study |
| pKK3 | pBG27 with *AlgE7*-Y122A | This study |
| pRS17 | pBG27 with *AlgE7*-Y122F | This study |
| pTB95 | pBG27 with *AlgE7*-R148G | This study |
| pJR4 | pBG27 with *AlgE7*-R148K | This study |
| pJR3 | pBG27 with *AlgE7*-Y149A | This study |
| pRS18 | pBG27 with *AlgE7*-Y149F | This study |
| pRS35 | pBG27 with *AlgE7*-Y149H | This study |
| pRS19 | pBG27 with *AlgE7*-D152E | This study |
| pRS20 | pBG27 with *AlgE7*-D152N | This study |
| pRS22 | pBG27 with *AlgE7*-H154F | This study |
| pRS21 | pBG27 with *AlgE7*-H154Y | This study |
| pJR1 | pBG27 with *AlgE7*-K172L | This study |
| pJR2 | pBG27 with *AlgE7*-K172R | This study |
| pKK5 | pBG27 with *AlgE7*-D173A | This study |
| pRS23 | pBG27 with *AlgE7*-D178E | This study |
| pRS24 | pBG27 with *AlgE7*-D178N | This study |
| pRS25 | pBG27 with *AlgE7*-D178R | This study |
| pRS26 | pBG27 with *AlgE7*-R195L | This study |
| pKK7 | pBG27 with *AlgE7*-H196A | This study |
| pKK10 | pBG27 with *AlgE7*-E117K-R148G | This study |
| pKK11 | pBG27 with *AlgE7*-E117K-R148G-K172L | This study |
| pJR5 | pBG27 with *AlgE7*-R148K-K172L | This study |
| pJR6 | pBG27 with *AlgE7*-R148K-K172R | This study |
| pTB113 | pBG27 with *AlgE7*-E117K-Y122F-R148G | This study |
| pTB118 | pBG27 with *AlgE7*-E117K-Y122F-K172L | This study |
| pTB119 | pBG27 with *AlgE7*-E117K-Y122F-K172R | This study |
| pTB111 | pBG27 with *AlgE7*-E117K-Y122F-R148G-K172L | This study |
| pTB112 | pBG27 with *AlgE7*-E117K-Y122F-R148G-K172R | This study |
| pTB116 | pBG27 with *AlgE7*-E117K-Y122F-Y149A | This study |

Enzyme extracts of AlgE7 wild type and mutants used in the A230 lyase activity assay were produced in 96 well plates as previously described (27). AlgE7 variant proteins and the wild type AlgE7 were expressed and purified in two different ways. Enzymes were partially purified after expression in DH5α at 37 °C, by fast protein liquid chromatography on a HiTrap-Q column (GE Healthcare), using two purification buffers and a disruption buffer containing 20 mM MOPS (pH 6.9) supplemented with 2 mM CaCl_2_, 2 mM CaCl_2_ + 1 M NaCl, and 4 mM CaCl_2_, respectively. AlgE7 and R148G were purified completely, by expressing the proteins in the pTYB1 vector containing a C-terminal intein tag in T7 Express competent *E. coli* cells (New England Biolabs) and purifying enzyme extract with affinity chromatography through the IMPACT™-CN system (New England Biolabs). Enzymes were eluted in HEPES buffer (20 mM HEPES, 5 mM CaCl_2_, 500 mM NaCl, pH 6.9) and dialysed in 5 mM HEPES and 5 mM CaCl_2_ to remove DTT and NaCl from the purification buffer, before they were freeze dried for storing.

Protein concentrations were estimated by the Bio-Rad protein assay (Bio-Rad) (43), or from the absorbance at 280 nm using a NanoDrop spectrophotometer and extinction coefficients calculated with the ExPASy ProtParam online tool (44).

### Activity assays

Two assays were performed for the AlgE7 variants, the tritium assay and the A230 assay. Tritium assay was performed on partially purified enzymes by measuring the liberation of ^3^H from [5-^3^H]-mannuronan, essentially as described previously (31). The alginate substrate and product were precipitated and removed from the sample before measuring the liberated ^3^H. Incubation with excess enzyme or prolonged incubation with a lyase will prevent efficient precipitation of the substrate due to the resulting short alginate oligomers, leading to an overestimation of the activity. Thus, appropriate enzyme amount and incubation times were identified to optimize this assay. Reaction buffer contained 20 mM MOPS (pH 6.9), and 2mM CaCl_2_, and reactions were incubated at 37 °C. The A230 absorbance assay was performed for crude enzyme extracts of variants by adding 20 µL of extracts to total reactions of 250 µL, 1 mg/ml polyM, in 50 mM Tris-buffer with 2.5 mM CaCl_2_. Absorbance was measured in COSTAR UV microplates every five minutes for 18 hours at 25 °C in a Spectramax ABS Plus microplate reader.

For AlgE7 purified with the IMPACT™-CN protocol, the A230 assay was performed in 50 mM Tris-buffer with various concentrations of NaCl (0, 50, 150, and 300 mM), CaCl_2_ (0, 1, and 3 mM), pH (6, 7, and 8), and polyM substrate (0.125 and 1 mg/ml), with a constant enzyme concentration of 0.3 µM. The reactions were performed in COSTAR UV 96 well plates, measuring absorbance at 230 nm every third minute, for 7-18 hours (until an end point was reached) at 25 °C. All reaction mixes contained 0.16 mM CaCl_2_ originating from the enzyme buffer. Each combination was performed twice.

### Product patterns with NMR and HPAEC-PAD

0,25 % w/v (2.5 mg/ml) of polyM or polyMG were dissolved in 5 mM HEPES buffer and 2.5 mM CaCl_2_, pH 6.9. AlgE7 and R148G purified with the IMPACT-CN protocol were added to a 1:300 w/w ratio with substrate, to final concentrations of about 8.3 µg/ml. The reactions were left at 25 °C for different time intervals. Reactions were stopped by adding EDTA to a final concentration of 5 mM, heated at 90 °C for 15 minutes and subsequently freeze dried. The samples were then dissolved in 600 µL D_2_O with 20 µL 0.3 M triethylenetetraamine-hexaacetate (TTHA), pH 7, as a calcium chelator, and 2.5 μL 1% 3-(trimethylsilyl)-propionic-2,2,3,3-d_4_ acid sodium salt (TSP) as a reference. NMR acquisition was done on a BRUKER AVIIIHD 400 MHz equipped with 5 mm SmartProbe (Bruker Biospin AG, Fällenden, Switzerland) at the NV-NMR-Centre/Norwegian NMR Platform (NNP) at the Norwegian University of Science and Technology (NTNU), a 30 ° flip angle pulse, 64 scans and a spectral width of 10 ppm. All spectra were recorded at 90 °C to decrease sample viscosity for increased resolution, and to move the solvent peak away from the chemical shift region of alginate. Integrations of spectra were done in TopSpin, and peaks were assigned as described previously (12, 45, 46). Sequential parameters were calculated according to the procedure in Supporting Information.

HPAEC-PAD was performed for the last time points of the AlgE7 reaction on polyM and polyMG (60 h), using oligomeric standards created by degradation of polyM and polyMG by an M-lyase from *Haliotis tuberculata*, and degradation of polyG by the G-lyase AlyA from *Klebsiella pneumonia* as previously described (42, 47).

### Time-resolved NMR

Time resolved ^13^C NMR experiments following the epimerization reaction were performed as previously described (14, 30), using 10 mg/ml of ^13^C1 enriched polyM, oligoG and polyMG. AlgE7 and R148G purified with the IMPACT-CN protocol were added to final concentrations of 2.8‒2.9 µM. The buffer contained 10 mM MOPS pH 6.9, 75 mM NaCl, and 2.5 mM CaCl_2_, in 99.8% D_2_O. For the wild type, the same experiment was also performed with varying concentrations of NaCl (0, 50, 75, 150, and 300 mM) and calcium (0, 1, 2.5, and 3 mM). After completion of the time-resolved experiment, a ^13^C-^1^H HSQC spectrum was recorded for assignment of the products formed during the enzymatic reaction. The time resolved data was acquired on 25 °C using a Bruker Avance III HD 800 MHz spectrometer (Bruker Biospin AG, Fällenden, Switzerland) equipped with a 5-mm Z-gradient CP-TCI (H/C/N) cryogenic probe at the NV-NMR-Centre/NNP at NTNU. All NMR spectra were recorded using TopSpin version 3.5.7. and processed in TopSpin version 4.0.8. software (Bruker BioSpin).

## Data availability

All data is contained within this manuscript and the Supporting Information.

## Supporting information

This article contains supporting information. The Supporting Information section cites the following references: (42, 46).

## Acknowledgements

We thank Rannveig Skrede for constructing the pRSxx plasmids (x denotes plasmid numbering), Karoline Kongsrud for constructing the pKKx plasmids, Jan Riedl for constructing the pJRx plasmids, Randi Aune for performing lyase activity assays, Olav Aarstad for performing the HPAEC-PAD analysis, and Wenche Iren Strand for assisting in NMR-analyses of lyase products.

## Funding and additional information

This research was funded by the Research Council of Norway through grants 294946 (The Norwegian Seaweed Biorefinery Platform), 250875 (Alginate Epimerase), and 226244 (Norwegian NMR platform-NNP).

## Abbreviations

Δ: 4-deoxy-L-*erythro*-hex-4-enepyranosyluronate
AA1-3: Amino Acid 1-3
CAZy: Carbohydrate-Active enZYmes
DP_n_: Number average degree of polymerization
G: α-L-guluronic acid
HPAEC-PAD: High-Performance Anion-Exchange Chromatography with Pulsed Amperometric Detection
HSQC: Heteronuclear Single Quantum Coherence spectroscopy
M: β-D-mannuronic acid

