## Supporting Information for "Mechanistic basis for understanding the dual activities of the bifunctional *Azotobacter vinelandii* mannuronan C-5 epimerase and alginate lyase AlgE7"

Figure S1: HSQC-spectra
Figure S2: Time-resolved $^{13}$C NMR, AlgE7 and oligoG
Figure S3: $^{1}$H NMR, calculation of sequential parameters
Figure S4: $^{1}$H NMR-spectra
Figure S5: HPAEC-PAD chromatograms
Figure S6: Protein sequences
Figure S7: Curves from factorial A230 lyase activity assay
Figure S8: End point values from AlgE7 factorial A230 lyase activity assay

Table S1: Fraction of signals from $^{1}$H NMR of AlgE7 and R148G at different time points
Table S2: Fraction of signals from $^{1}$H NMR of variants
Table S3: Slopes and end point values from AlgE7 factorial A230 lyase activity assay

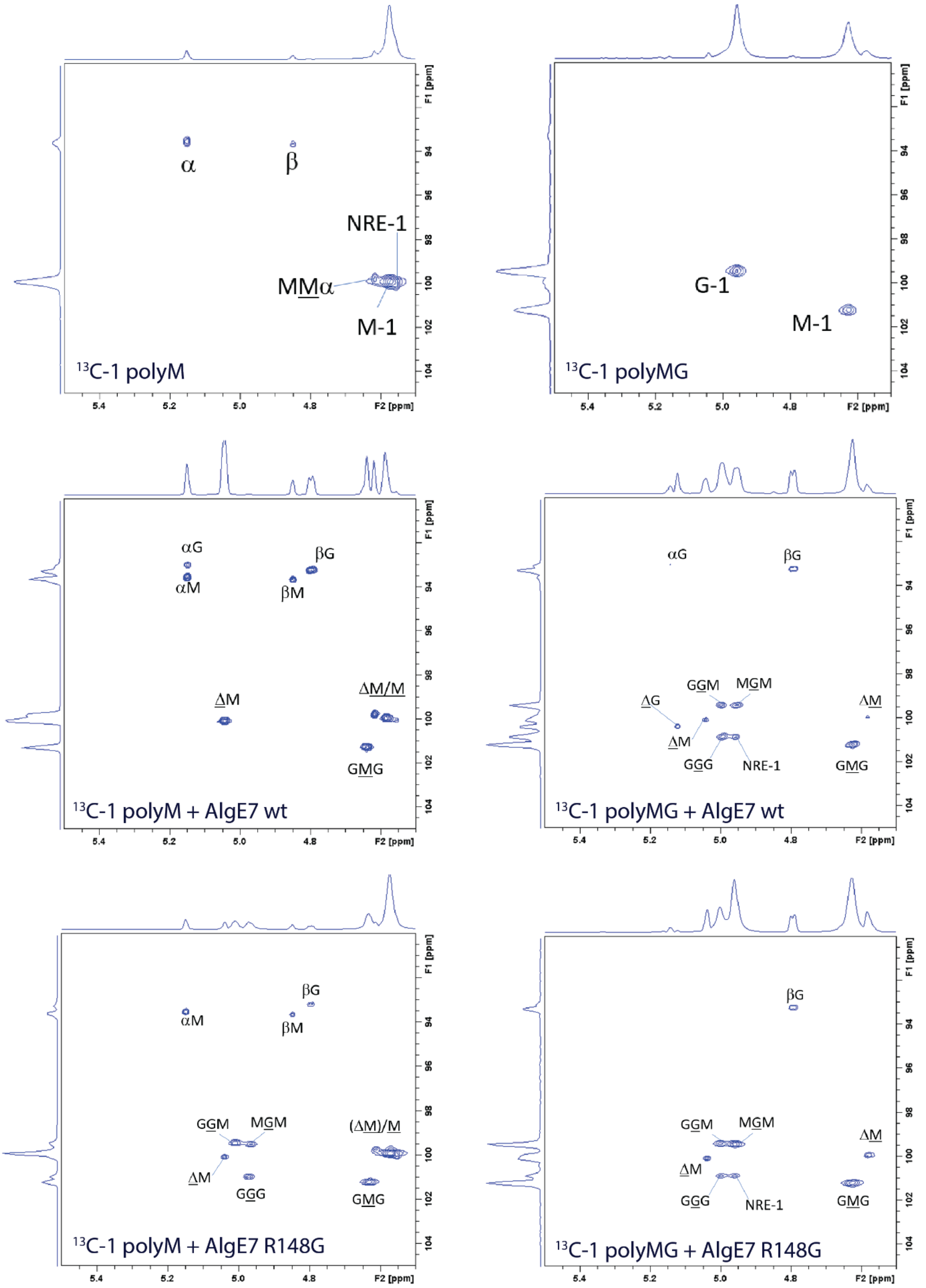


**Figure S1:** HSQC spectra of reaction products from 2.8–2.9 µM AlgE7 wild type and R148G on 10 mg/ml polyM or polyMG, in buffer containing 75 mM NaCl and 2.5 mM NaCl, pH 7.0. The spectra were recorded at the end of time-resolved $^{13}$C-NMR experiments, and assigned as shown, according to previous work (Grasdalen et al. 1981; Grasdalen 1983; Holtan, Zhang, et al. 2006). Triads or dyads are indicated in the spectra, and the residue giving rise to the signal is underlined. NRE-1 denotes the signal from carbon 1 at the non-reducing end.

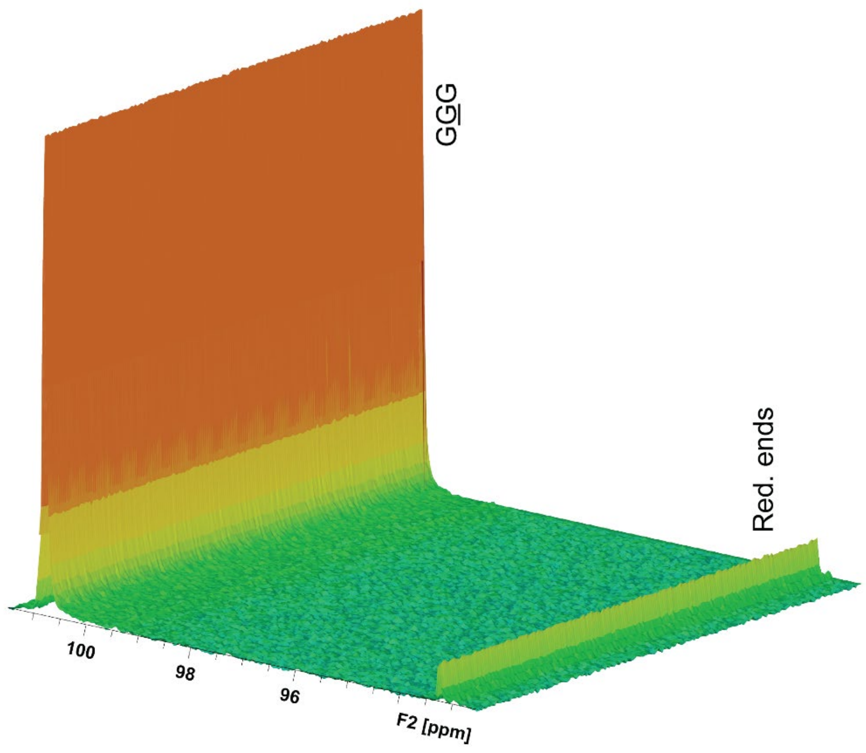


**Figure S2:** OligoG substrate incubated with AlgE7 produced no change in $^{13}C$ time resolved NMR spectra after 16 hours. The same test was performed with R148G, and as the results were identical only the spectrum for the wild type is shown.

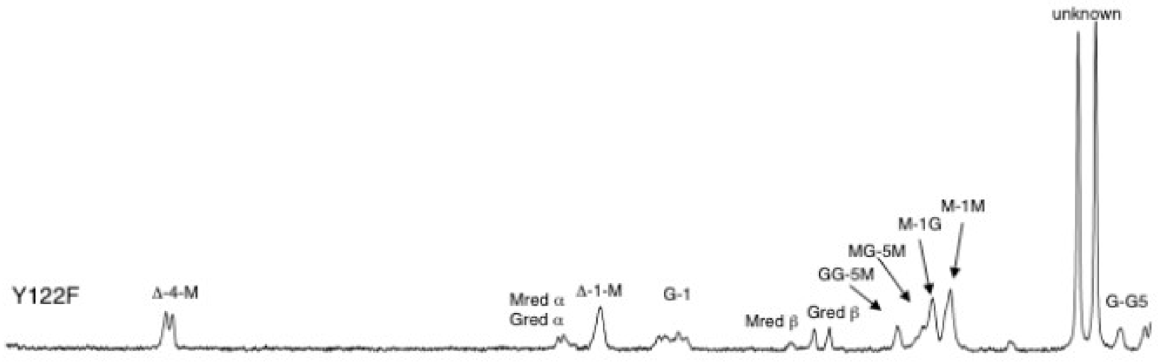


**Figure S3:** Example chromatogram showing the anomeric region of polyM epimerized by the AlgE7 variant Y122F. The signal marked "unknown" appears in all NMR-spectra from enzyme extracts, and does not appear when purified enzymes are used. The G-1 peak comes from all internal G-residues, but for some of the active lyases it almost disappeared. However, there were clearly internal G-residues present, seen from the peaks of GGM-5, MGM-5 and MG-1. Due to poor baseline separation, the peaks for MGM-5 and MG-1 were difficult to integrate separately. Nevertheless, this was done, as the two peaks contain some of the information required to calculate the amount of total G- and M-residues in the sample. The results are not as robust as they would be for the maximum averaging procedure that is possible to do for epimerized alginate with high $DP_n$, but they can still be used for qualitative comparisons of different time points and variants. Calculations of sequential parameters of epimerized and lyased alginate were done using the following relationships between signal intensities:

1. I(total) = I(GGM-5) + I(MGM-5) + I(MG-1) + I(MM-1) + I(GG-5) + I(Δ) + $I(M_{red})$ + $I(G_{red})$
2. $F(G_{int})$ = I(G-1)/I(total)
3. F(GGM) = I(GGM-5) / I(total)
4. F(MM) = I(MM-1) / I(total)
5. F(GG) = I(GG-5) / I(total)
6. F(Δ) = I(ΔM-1) / I(total)
7. $F(G_{red}) = (I(G_{red}\beta) + I((G_{red}\beta) \cdot 0.2)) / I(total)$
8. $F(M_{red}) = (M_{red}\beta + (M_{red}\beta \cdot 2.2)) / I(total)$

9. F(G) = ( I(GGM-5) + I(MGM-5) + I(GG-5) + $I(G_{red})$ ) / I(total)
10. F(M) = ( I(MG-1) + I(MM-1) + $I(M_{red})$ ) / I(total)
11. $F(M_{int}) = F(M)-F(M_{red})$

As the signals for M and G reducing ends in α-configuration are not resolved, the known relationships of $G_{red}\alpha$ being 0.2 of $G_{red}\beta$, and $M_{red}\alpha$ being 2.2 times more than $M_{red}\beta$, are used (Heyraud et al. 1996).

Fractions were calculated by dividing the relevant intensity by the total intensity of all peaks. Number average degree of polymerization ($DP_n$) was calculated as following:

$$DP_n = \frac{I(\text{total})}{I(M_{red}) + I(G_{red})}$$

Based on these calculations, we obtained the values listed in Table S2 from incubating partially purified protein extracts of AlgE7 wild type and variants with 1 mg/ml polyM for 24 hours. In Table S1 the data for purified wild type AlgE7 and R148G incubated with 2.5 mg/ml substrate (enzyme:substrate ratio 1:300, w/w) at different reaction time points is listed.

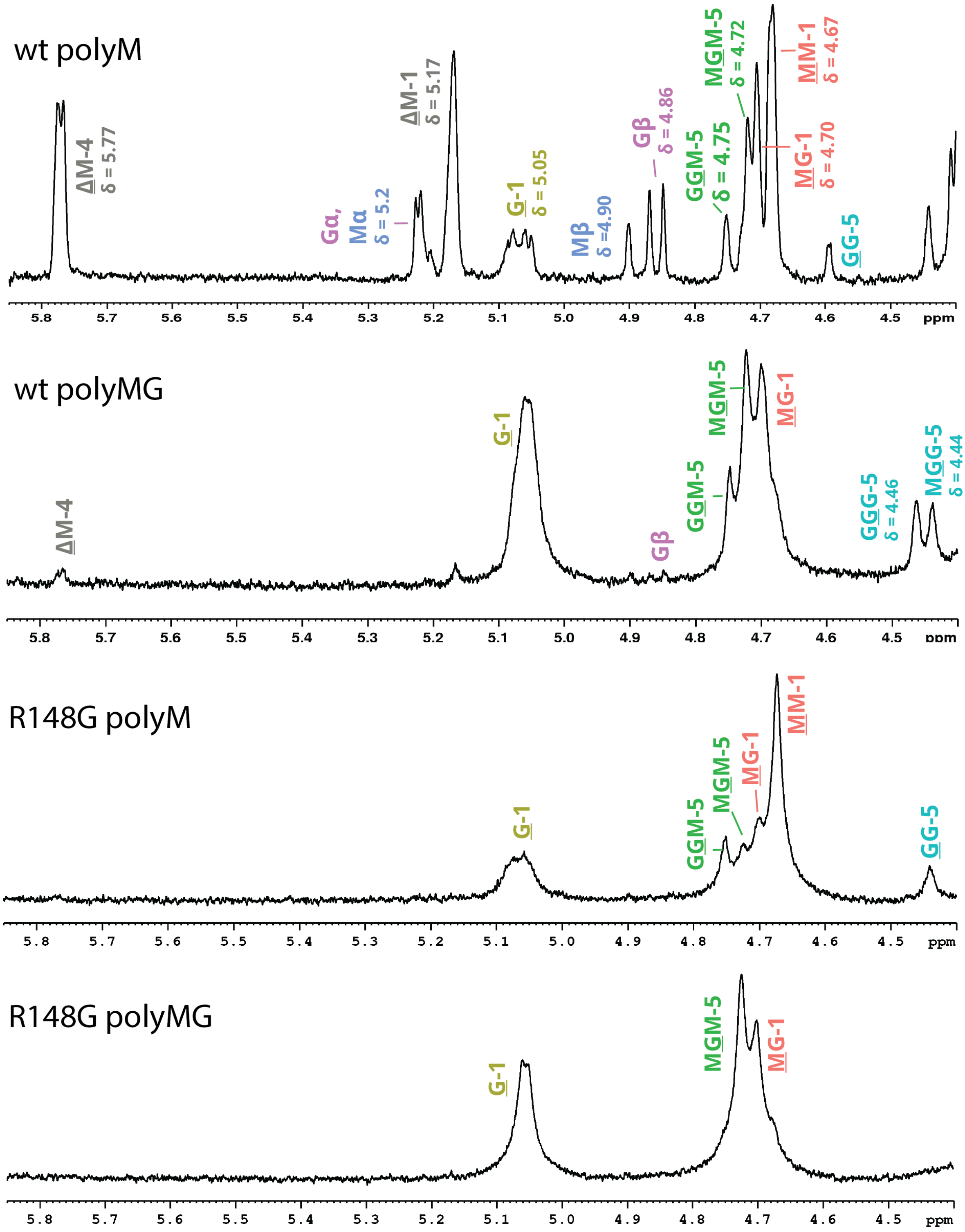


**Figure S4**: $^{1}$H-NMR of reactions of AlgE7 (wt) and R148G on polyM and polyMG, with no added NaCl, after 60 h of enzyme reaction. Compare with Figures 2 and 5 for the results with 75 mM NaCl added.

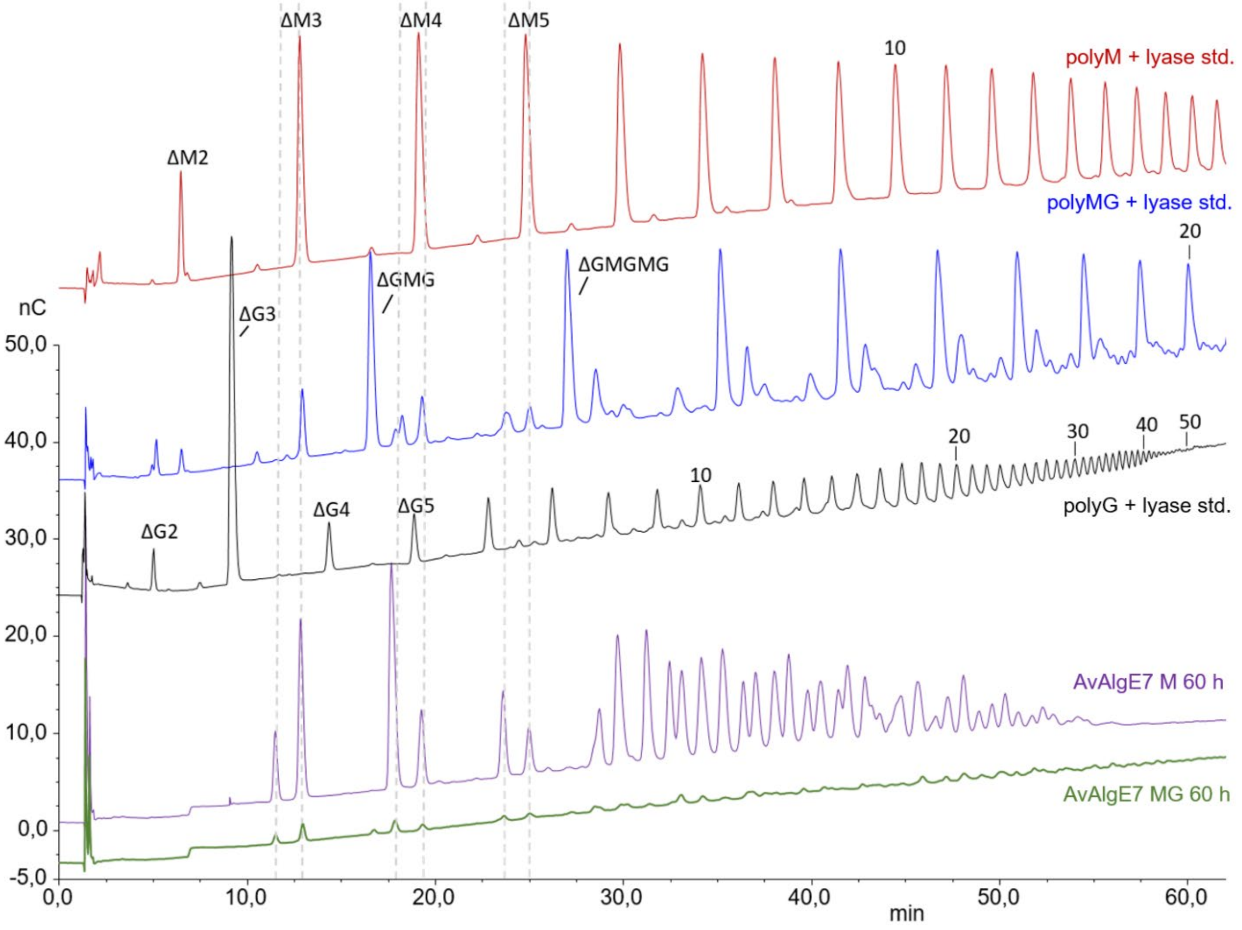


**Figure S5**: HPAEC-PAD chromatogram of the products produced by AlgE7 after incubation of 60 h on polyM (purple) and polyMG (green). Standards of M-lyase from *Haliotis tubercalata* on polyM and polyMG, and G-lyase AlyA from *Klebsiella pneumoniae* on polyG, are shown in red, blue and black lines, respectively. Relevant, annotated peaks of the standards are marked (Aarstad et al. 2012).

```
                10         20         30         40         50         60         70         80         90        100
AlgE7  MEYNVKDFGAKGDGKTDDTDAIQAAIDAAHKAGGGTVYLPSGEYRVSGGDEASDGALIIKSNVYIVGAGMGETVIKLVDGWDEKLTGIIRSANGEKTHDY
ConE   .DX.......L...VS..XA.........XA.........A......XXX.P...C.XX.X...X...............X.QXX..MV...Y..E.SNF
ConL   ..F.........................X.........A...............M......M................Q........K........

               110        120        130        140        150        160        170        180        190        200
AlgE7  GISDLTIDGNQDNTEGEVDGFYTGYIPGKNGADYNVTVERVEIREVSRYAFDPHEQTINLTIRDSVAHDNGKDGFVADFQIGAVFENNVSYNNGRHGFNI
ConE   .M....L...R...SXK...WFN.....XD...RD..L.......M.G.G..................L......YXXXX......X...D......
ConL   ......L...................X.D.................................................DST...............

               210        220        230        240        250        260        270        280        290        300
AlgE7  VTSSHDIVFTNNVAYGNGANGLVVQRGSEDRDFVYNVEIEGGSFHDNGQEGVLIKMSTDVTLQGAEIYGNGYAGVRVQGVEDVRILDNYIHDNAQSKANA
ConE   ...TX.F.XX........GA........X.LAHPXXIL.D..AYY..XL...XX..XX.....N.......XX....Y.AX..Q....Q....X.NGXYX
ConL   .........R.....................XP..IQ....AY....X....................A.........X.Q...........XG...

               310        320        330          340         350          360        370           380
AlgE7  EVIVESYDDRDGPSDDYYETQNVTVKGNTIVGSAN--STYGIQ--ERADGTDYTSI--GNNSVSGTQRGIVQLSGTNSTFS----GRSGDAYQFIDGST
ConE   ..LLQ....TA.V.GNF.X.XXTWIE..X.X....--......--..X.....S.L--YA.XIX.V.X.X.X.Y.A..XV.----XXX.XXG.QAT---
ConL   .IXL.X......X.GN....L.A..Q.........XX....XXXL.....X..XXXXX.X..X.......X.X...XX...XXXXX.FA...XX.XXXXX
```

**Figure S6:** Sequence alignment of AlgE7 and consensus sequences of the nine A-modules that are pure epimerases (ConE) and the three A-modules that are bifunctional lyases/epimerases (conL) from *A. vinelandii* and *A. chroococcum*. The active site residues are highlighted in purple, and residues mutated in this study are highlighted in blue. An excerpt of this is also shown in Figure 3A.

wt, 1 mg/ml, pH 7

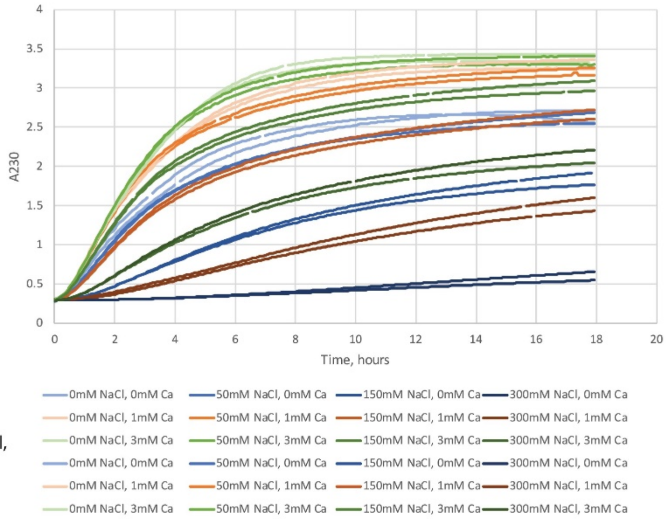


wt, 0.125 mg/ml, pH 7

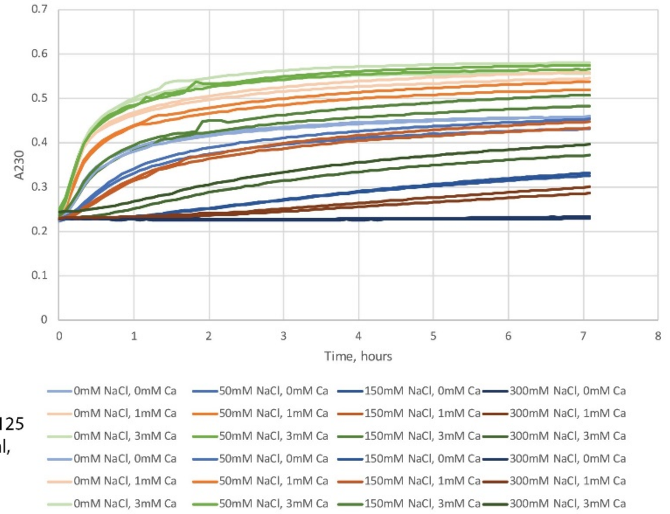


**Figure S7:** Lyase activity over time measured from the spectrophotometric A230 assay for AlgE7 on two different concentrations of polyM and different concentrations of NaCl and $CaCl_2$. Similar curves were obtained at pH 6 and 8, and for the amino acid replacement variants. Initial, linear regions of the curves were used to calculate the slopes, listed in Table 1 as percentages of wild type for the enzyme variants, and in Table S3 for the factorial assay together with values at end point (with subtracted start values). The slopes are also presented in Figure 6, and the values at end point are shown in Figure S8.

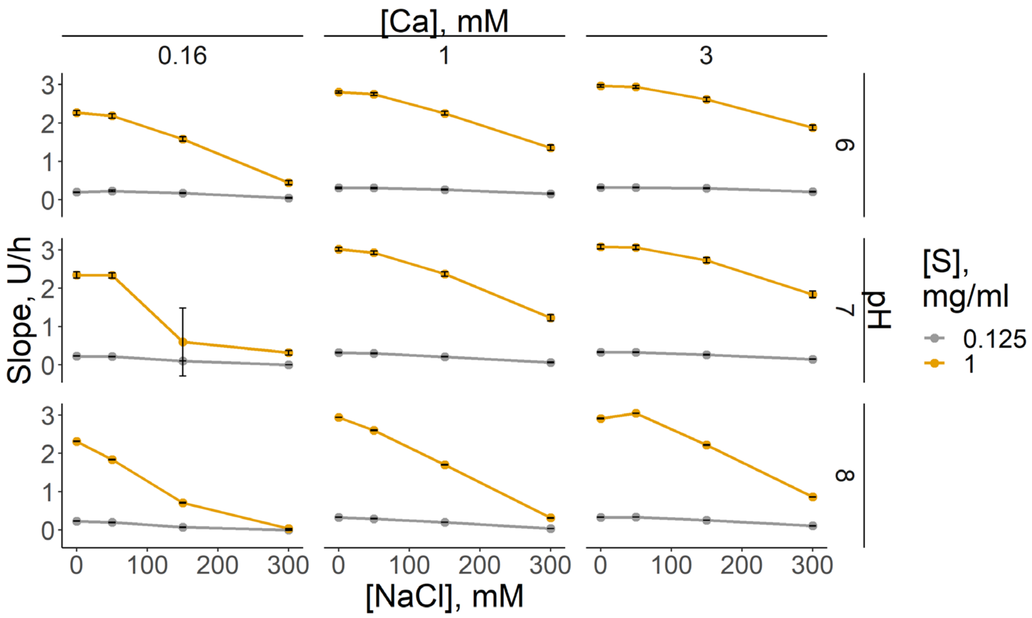


**Figure S8:** Mean end point values of factorial lyase assay of the wild type AlgE7. Points are averages of two repetitions, and error bars show the range of values for the repetitions.

**Table S1**: Fraction of signals from end point $^{1}$H NMR of wild type AlgE7 and R148G at different reaction time points, analysed with $^{1}$H NMR.

| Sample | Time (h) | F(G) | F(M) | F(GG) | F(MM) | $F(G_{int})$ | $F(M_{int})$ | F(GGM) | F(Δ) | $F(G_{red})$ | $F(M_{red})$ | DPn |
|---|---|---|---|---|---|---|---|---|---|---|---|---|
| Wt + polyM | 0 | 0.00 | 1.00 | 0.00 | 1.00 | 0.00 | 1.00 | 0.00 | 0.00 | 0.00 | 0.00 | NaN |
| 0 mM NaCl | 6 | 0.18 | 0.80 | 0.04 | 0.67 | 0.12 | 0.80 | 0.04 | 0.02 | 0.01 | 0.01 | 78.09 |
| | 12 | 0.22 | 0.74 | 0.05 | 0.53 | 0.18 | 0.70 | 0.06 | 0.04 | 0.02 | 0.04 | 17.18 |
| | 24 | 0.25 | 0.65 | 0.07 | 0.38 | 0.19 | 0.58 | 0.07 | 0.10 | 0.05 | 0.06 | 9.19 |
| | 48 | 0.30 | 0.55 | 0.09 | 0.31 | 0.16 | 0.49 | 0.06 | 0.15 | 0.06 | 0.07 | 7.89 |
| | 60 | 0.30 | 0.50 | 0.06 | 0.26 | 0.11 | 0.41 | 0.04 | 0.20 | 0.09 | 0.09 | 5.56 |
| Wt + polyMG | 0 | 0.49 | 0.51 | 0.03 | 0.12 | 0.49 | 0.51 | 0.00 | 0.00 | 0.00 | 0.00 | NaN |
| 0 mM NaCl | 6 | 0.57 | 0.42 | 0.03 | 0.00 | 0.53 | 0.40 | 0.13 | 0.01 | 0.01 | 0.01 | NaN |
| | 12 | 0.53 | 0.47 | 0.05 | 0.00 | 0.56 | 0.45 | 0.11 | 0.01 | 0.01 | 0.02 | NaN |
| | 24 | 0.58 | 0.40 | 0.09 | 0.00 | 0.59 | 0.39 | 0.14 | 0.02 | 0.01 | 0.01 | 54.81 |
| | 48 | 0.58 | 0.40 | 0.13 | 0.00 | 0.60 | 0.39 | 0.14 | 0.02 | 0.01 | 0.01 | 44.85 |
| | 60 | 0.61 | 0.37 | 0.16 | 0.00 | 0.65 | 0.35 | 0.14 | 0.02 | 0.01 | 0.02 | 29.98 |
| R148G + polyM | 0 | 0.00 | 1.00 | 0.00 | 1.00 | 0.00 | 1.00 | 0.00 | 0.00 | 0.00 | 0.00 | NaN |
| 0 mM NaCl | 6 | 0.17 | 0.83 | 0.01 | 0.71 | 0.07 | 0.80 | 0.06 | 0.01 | 0.02 | 0.02 | NaN |
| | 12 | 0.18 | 0.82 | 0.02 | 0.69 | 0.10 | 0.82 | 0.07 | 0.00 | 0.01 | 0.01 | 59.50 |
| | 24 | 0.22 | 0.77 | 0.04 | 0.62 | 0.16 | 0.76 | 0.08 | 0.01 | 0.01 | 0.02 | 40.40 |
| | 48 | 0.27 | 0.71 | 0.05 | 0.54 | 0.21 | 0.70 | 0.12 | 0.01 | 0.01 | 0.01 | 43.25 |
| | 60 | 0.27 | 0.72 | 0.06 | 0.53 | 0.24 | 0.70 | 0.11 | 0.01 | 0.01 | 0.02 | 33.06 |
| R148G + polyMG | 0 | 0.49 | 0.51 | 0.00 | 0.12 | 0.49 | 0.51 | 0.00 | 0.00 | 0.00 | 0.00 | NaN |
| 0 mM NaCl | 6 | 0.56 | 0.44 | 0.04 | 0.12 | 0.48 | 0.44 | 0.10 | 0.00 | 0.00 | 0.00 | NaN |
| | 12 | 0.53 | 0.47 | 0.03 | 0.14 | 0.48 | 0.47 | 0.10 | 0.00 | 0.00 | 0.00 | NaN |
| | 24 | 0.54 | 0.46 | 0.03 | 0.13 | 0.48 | 0.46 | 0.10 | 0.00 | 0.00 | 0.00 | NaN |
| | 48 | 0.54 | 0.46 | 0.03 | 0.13 | 0.48 | 0.46 | 0.10 | 0.00 | 0.00 | 0.00 | NaN |
| | 60 | 0.55 | 0.45 | 0.03 | 0.12 | 0.49 | 0.45 | 0.10 | 0.00 | 0.00 | 0.00 | NaN |
| Wt + polyM | 0 | 0.00 | 1.00 | 0.00 | 1.00 | 0.00 | 1.00 | 0.00 | 0.00 | 0.00 | 0.00 | NaN |
| 75 mM NaCl | 6 | 0.15 | 0.84 | 0.03 | 0.72 | 0.09 | 0.84 | 0.04 | 0.01 | 0.00 | 0.00 | 352.53 |
| | 12 | 0.20 | 0.78 | 0.06 | 0.63 | 0.14 | 0.77 | 0.05 | 0.02 | 0.01 | 0.01 | 71.61 |
| | 24 | 0.23 | 0.71 | 0.07 | 0.50 | 0.19 | 0.68 | 0.07 | 0.06 | 0.02 | 0.02 | 21.74 |
| | 48 | 0.28 | 0.60 | 0.09 | 0.35 | 0.16 | 0.55 | 0.07 | 0.12 | 0.05 | 0.05 | 10.07 |
| | 60 | 0.28 | 0.58 | 0.07 | 0.32 | 0.17 | 0.51 | 0.06 | 0.14 | 0.07 | 0.06 | 7.62 |
| Wt + polyMG | 0 | 0.54 | 0.46 | 0.03 | 0.12 | 0.49 | 0.46 | 0.00 | 0.00 | 0.00 | 0.00 | NaN |
| 75 mM NaCl | 6 | 0.61 | 0.41 | 0.10 | 0.08 | 0.55 | 0.41 | 0.10 | 0.02 | 0.01 | 0.00 | 92.02 |
| | 12 | 0.54 | 0.44 | 0.12 | 0.08 | 0.58 | 0.43 | 0.14 | 0.03 | 0.02 | 0.01 | 38.16 |
| | 24 | 0.56 | 0.41 | 0.16 | 0.06 | 0.61 | 0.40 | 0.15 | 0.03 | 0.02 | 0.00 | 51.86 |
| | 48 | 0.57 | 0.40 | 0.21 | 0.06 | 0.63 | 0.39 | 0.15 | 0.04 | 0.02 | 0.01 | 32.37 |
| | 60 | 0.59 | 0.37 | 0.22 | 0.05 | 0.64 | 0.37 | 0.16 | 0.04 | 0.02 | 0.01 | 41.26 |
| R148G + polyM | 0 | 0.00 | 1.00 | 0.00 | 1.00 | 0.00 | 1.00 | 0.00 | 0.00 | 0.00 | 0.00 | NaN |
| 75 mM NaCl | 6 | 0.11 | 0.89 | -0.02 | 0.83 | 0.03 | 0.89 | 0.05 | 0.00 | 0.00 | 0.00 | NaN |
| | 12 | 0.14 | 0.85 | -0.02 | 0.73 | 0.07 | 0.85 | 0.07 | 0.00 | 0.00 | 0.00 | NaN |
| | 24 | 0.18 | 0.82 | 0.00 | 0.70 | 0.09 | 0.82 | 0.08 | 0.00 | 0.00 | 0.00 | NaN |
| | 48 | 0.22 | 0.76 | 0.04 | 0.61 | 0.18 | 0.75 | 0.10 | 0.01 | 0.01 | 0.02 | 33.34 |
| | 60 | 0.25 | 0.75 | 0.05 | 0.61 | 0.17 | 0.75 | 0.10 | 0.00 | 0.00 | 0.00 | NaN |

| | | | | | | | | | | | | |
|---|---|---|---|---|---|---|---|---|---|---|---|---|
| R148G + polyMG | 0 | 0.49 | 0.51 | 0.00 | 0.12 | 0.49 | 0.51 | 0.00 | 0.00 | 0.00 | 0.00 | NaN |
| 75 mM NaCl | 6 | 0.56 | 0.45 | 0.03 | 0.09 | 0.50 | 0.45 | 0.07 | 0.00 | 0.00 | 0.00 | NaN |
| | 12 | 0.53 | 0.48 | 0.04 | 0.14 | 0.48 | 0.48 | 0.08 | 0.00 | 0.00 | 0.00 | NaN |
| | 24 | 0.58 | 0.42 | 0.05 | 0.10 | 0.51 | 0.42 | 0.09 | 0.02 | 0.00 | 0.00 | NaN |
| | 48 | 0.59 | 0.42 | 0.06 | 0.10 | 0.52 | 0.42 | 0.10 | 0.01 | 0.01 | 0.00 | 82.38 |
| | 60 | 0.59 | 0.42 | 0.07 | 0.10 | 0.52 | 0.42 | 0.10 | 0.02 | 0.01 | 0.00 | 80.84 |

**Table S2:** Fraction of signals from end point $^{1}$H NMR of all variants tested in this study. $G_{red}$ and $M_{red}$ are residues at reducing ends, and $G_{int}$ are internal G-residues.

| Enzyme | F(G) | F(M) | F(GG) | F(MM) | F($G_{int}$) | F($M_{int}$) | F(GGM) | F(Δ) | F($G_{red}$) | F($M_{red}$) | DPn |
|---|---|---|---|---|---|---|---|---|---|---|---|
| Wt | 0.24 | 0.47 | 0.01 | 0.22 | 0.13 | 0.34 | 0.00 | 0.29 | 0.11 | 0.14 | 4.02 |
| Wt | 0.25 | 0.47 | 0.01 | 0.21 | 0.14 | 0.32 | 0.01 | 0.28 | 0.11 | 0.15 | 3.86 |
| E117L | 0.30 | 0.45 | 0.06 | 0.23 | 0.20 | 0.35 | 0.02 | 0.25 | 0.11 | 0.09 | 4.94 |
| E117L | 0.30 | 0.49 | 0.05 | 0.25 | 0.20 | 0.40 | 0.04 | 0.21 | 0.10 | 0.09 | 5.39 |
| E117K | 0.31 | 0.52 | 0.07 | 0.26 | 0.21 | 0.43 | 0.07 | 0.17 | 0.10 | 0.09 | 5.17 |
| E117K | 0.30 | 0.49 | 0.05 | 0.26 | 0.19 | 0.40 | 0.04 | 0.20 | 0.11 | 0.09 | 4.96 |
| D119A | 0.00 | 1.00 | 0.00 | 1.00 | 0.00 | 1.00 | 0.00 | 0.00 | 0.00 | 0.00 | NaN |
| D119A | 0.00 | 1.00 | 0.00 | 1.00 | 0.00 | 1.00 | 0.00 | 0.00 | 0.00 | 0.00 | NaN |
| D119E | 0.26 | 0.62 | 0.08 | 0.39 | 0.21 | 0.56 | 0.07 | 0.11 | 0.05 | 0.06 | 9.18 |
| D119E | 0.26 | 0.67 | 0.07 | 0.48 | 0.22 | 0.64 | 0.07 | 0.07 | 0.04 | 0.03 | 14.10 |
| D119N | 0.00 | 1.00 | 0.00 | 1.00 | 0.00 | 1.00 | 0.00 | 0.00 | 0.00 | 0.00 | NaN |
| D119N | 0.00 | 1.00 | 0.00 | 1.00 | 0.00 | 1.00 | 0.00 | 0.00 | 0.00 | 0.00 | NaN |
| Y122A | 0.00 | 1.00 | 0.00 | 1.00 | 0.00 | 1.00 | 0.00 | 0.00 | 0.00 | 0.00 | NaN |
| Y122A | 0.00 | 1.00 | 0.00 | 1.00 | 0.00 | 1.00 | 0.00 | 0.00 | 0.00 | 0.00 | NaN |
| Y122F | 0.32 | 0.51 | 0.08 | 0.26 | 0.22 | 0.45 | 0.07 | 0.17 | 0.10 | 0.05 | 6.54 |
| Y122F | 0.32 | 0.50 | 0.06 | 0.24 | 0.21 | 0.42 | 0.06 | 0.18 | 0.11 | 0.08 | 5.25 |
| R148G | 0.20 | 0.80 | 0.02 | 0.69 | 0.18 | 0.79 | 0.07 | 0.00 | 0.01 | 0.01 | 51.60 |
| R148G | 0.31 | 0.67 | 0.07 | 0.51 | 0.30 | 0.66 | 0.12 | 0.02 | 0.01 | 0.01 | 45.41 |
| R148K | 0.34 | 0.62 | 0.13 | 0.42 | 0.32 | 0.60 | 0.11 | 0.04 | 0.02 | 0.02 | 25.61 |
| R148K | 0.30 | 0.66 | 0.08 | 0.46 | 0.27 | 0.64 | 0.10 | 0.04 | 0.03 | 0.02 | 21.62 |
| K172L | 0.29 | 0.49 | 0.03 | 0.25 | 0.18 | 0.39 | 0.03 | 0.22 | 0.11 | 0.09 | 4.95 |
| K172L | 0.37 | 0.47 | 0.12 | 0.24 | 0.26 | 0.39 | 0.05 | 0.17 | 0.11 | 0.08 | 5.29 |
| K172R | 0.25 | 0.46 | 0.00 | 0.21 | 0.13 | 0.32 | 0.00 | 0.29 | 0.12 | 0.14 | 3.90 |
| K172R | 0.24 | 0.47 | 0.01 | 0.22 | 0.12 | 0.33 | 0.00 | 0.29 | 0.12 | 0.13 | 3.93 |
| D173A | 0.00 | 1.00 | 0.00 | 1.00 | 0.00 | 1.00 | 0.00 | 0.00 | 0.00 | 0.00 | NaN |
| D173A | 0.00 | 1.00 | 0.00 | 1.00 | 0.00 | 1.00 | 0.00 | 0.00 | 0.00 | 0.00 | NaN |
| E117K + R148G | 0.36 | 0.61 | 0.13 | 0.40 | 0.35 | 0.60 | 0.13 | 0.02 | 0.01 | 0.02 | 35.38 |
| E117K + R148G | 0.34 | 0.65 | 0.08 | 0.47 | 0.32 | 0.64 | 0.13 | 0.01 | 0.02 | 0.01 | 29.45 |
| E117K + R148G + K127L | 0.39 | 0.58 | 0.12 | 0.37 | 0.37 | 0.56 | 0.16 | 0.03 | 0.03 | 0.02 | 20.23 |
| E117K + R148G + K127L | 0.38 | 0.60 | 0.12 | 0.39 | 0.36 | 0.58 | 0.15 | 0.02 | 0.02 | 0.02 | 26.57 |
| R148G + K172L | 0.25 | 0.75 | 0.08 | 0.62 | 0.25 | 0.75 | 0.09 | 0.00 | 0.00 | 0.00 | NaN |
| R148G + K172L | 0.19 | 0.81 | 0.04 | 0.68 | 0.19 | 0.81 | 0.08 | 0.00 | 0.00 | 0.00 | NaN |
| R148G + K172R | 0.32 | 0.66 | 0.06 | 0.48 | 0.31 | 0.65 | 0.13 | 0.02 | 0.01 | 0.01 | 49.25 |
| R148G + K172R | 0.31 | 0.68 | 0.08 | 0.51 | 0.30 | 0.68 | 0.12 | 0.01 | 0.01 | 0.00 | 140.97 |
| E117K + Y122F + R148G | 0.40 | 0.59 | 0.15 | 0.38 | 0.40 | 0.59 | 0.17 | 0.01 | 0.00 | 0.00 | NaN |
| E117K + Y122F + R148G | 0.39 | 0.60 | 0.11 | 0.41 | 0.38 | 0.60 | 0.17 | 0.01 | 0.01 | 0.00 | 85.36 |
| E117K + Y122F + K172L | 0.39 | 0.50 | 0.16 | 0.25 | 0.34 | 0.49 | 0.14 | 0.11 | 0.05 | 0.00 | 18.96 |
| E117K + Y122F + K172L | 0.39 | 0.50 | 0.16 | 0.25 | 0.33 | 0.49 | 0.13 | 0.11 | 0.06 | 0.01 | 13.85 |
| E117K + Y122F + K172R | 0.39 | 0.54 | 0.15 | 0.29 | 0.35 | 0.50 | 0.13 | 0.07 | 0.04 | 0.04 | 12.56 |
| E117K + Y122F + K172R | 0.38 | 0.57 | 0.14 | 0.34 | 0.34 | 0.54 | 0.14 | 0.06 | 0.04 | 0.02 | 17.20 |
| E117K + Y122F + R148G + K172L | 0.45 | 0.54 | 0.21 | 0.35 | 0.45 | 0.54 | 0.15 | 0.01 | 0.00 | 0.00 | NaN |
| E117K + Y122F + R148G + K172L | 0.35 | 0.63 | 0.11 | 0.45 | 0.35 | 0.63 | 0.15 | 0.02 | 0.00 | 0.00 | NaN |

| | | | | | | | | | | | |
|---|---|---|---|---|---|---|---|---|---|---|---|
| E117K + Y122F + R148G + K172R | 0.33 | 0.65 | 0.11 | 0.50 | 0.31 | 0.64 | 0.12 | 0.02 | 0.01 | 0.01 | 46.73 |
| E117K + Y122F + R148G + K172R | 0.28 | 0.71 | 0.10 | 0.56 | 0.28 | 0.71 | 0.10 | 0.01 | 0.00 | 0.00 | NaN |

**Table S3:** Calculated slopes and end point values (initial value subtracted) for all the lyase activity experiments performed for wild type AlgE7 at three different pH values, two different substrate concentrations, four different salt concentrations, and three different calcium concentrations. Each experiment was repeated twice, and values from both experiments are shown.

| | mM | | 1 mg/ml [S] | | | | 0.125 mg/ml ]S] | | | |
|---|---|---|---|---|---|---|---|---|---|---|
| pH | [NaCl] | [Ca] | Slope | | End value | | Slope | | End value | |
| 6 | 0 | 0 | 0.25 | 0.24 | 2.49 | 2.61 | 0.22 | 0.20 | 0.42 | 0.44 |
| | 50 | 0 | 0.24 | 0.25 | 2.41 | 2.54 | 0.16 | 0.17 | 0.45 | 0.48 |
| | 150 | 0 | 0.09 | 0.09 | 1.80 | 1.92 | 0.00 | 0.01 | 0.39 | 0.42 |
| | 300 | 0 | 0.00 | 0.01 | 0.67 | 0.78 | 0.00 | 0.00 | 0.27 | 0.29 |
| | 0 | 1 | 0.32 | 0.32 | 3.06 | 3.11 | 0.30 | 0.32 | 0.53 | 0.56 |
| | 50 | 1 | 0.34 | 0.34 | 3.00 | 3.07 | 0.33 | 0.32 | 0.53 | 0.56 |
| | 150 | 1 | 0.25 | 0.25 | 2.49 | 2.59 | 0.11 | 0.11 | 0.49 | 0.50 |
| | 300 | 1 | 0.04 | 0.05 | 1.57 | 1.71 | 0.01 | 0.01 | 0.37 | 0.41 |
| | 0 | 3 | 0.35 | 0.34 | 3.24 | 3.30 | 0.32 | 0.33 | 0.54 | 0.57 |
| | 50 | 3 | 0.39 | 0.39 | 3.21 | 3.29 | 0.37 | 0.38 | 0.54 | 0.57 |
| | 150 | 3 | 0.35 | 0.36 | 2.85 | 2.95 | 0.23 | 0.24 | 0.53 | 0.54 |
| | 300 | 3 | 0.14 | 0.15 | 2.10 | 2.23 | 0.02 | 0.01 | 0.43 | 0.45 |
| 7 | 0 | 0 | 0.55 | 0.48 | 2.55 | 2.70 | 0.24 | 0.23 | 0.46 | 0.46 |
| | 50 | 0 | 0.41 | 0.40 | 2.55 | 2.68 | 0.09 | 0.09 | 0.43 | 0.45 |
| | 150 | 0 | 0.07 | 0.07 | 1.77 | 0.00 | 0.00 | 0.00 | 0.33 | 0.33 |
| | 300 | 0 | 0.00 | 0.01 | 0.55 | 0.66 | 0.00 | 0.00 | 0.23 | 0.23 |
| | 0 | 1 | 0.65 | 0.67 | 3.25 | 3.36 | 0.49 | 0.50 | 0.54 | 0.56 |
| | 50 | 1 | 0.61 | 0.63 | 3.17 | 3.26 | 0.40 | 0.41 | 0.52 | 0.54 |
| | 150 | 1 | 0.33 | 0.34 | 2.61 | 2.72 | 0.06 | 0.06 | 0.43 | 0.45 |
| | 300 | 1 | 0.03 | 0.04 | 1.43 | 1.60 | 0.00 | 0.00 | 0.29 | 0.30 |
| | 0 | 3 | 0.63 | 0.64 | 3.32 | 3.43 | 0.51 | 0.52 | 0.56 | 0.58 |
| | 50 | 3 | 0.62 | 0.63 | 3.30 | 3.41 | 0.49 | 0.51 | 0.57 | 0.57 |
| | 150 | 3 | 0.50 | 0.53 | 2.96 | 3.09 | 0.22 | 0.22 | 0.48 | 0.51 |
| | 300 | 3 | 0.13 | 0.14 | 2.04 | 2.21 | 0.01 | 0.01 | 0.37 | 0.40 |
| 8 | 0 | 0 | 0.39 | 0.38 | 2.61 | 2.60 | 0.14 | 0.15 | 0.47 | 0.48 |
| | 50 | 0 | 0.17 | 0.16 | 2.14 | 2.12 | 0.00 | 0.01 | 0.44 | 0.44 |
| | 150 | 0 | 0.01 | 0.01 | 1.00 | 0.99 | 0.01 | 0.00 | 0.32 | 0.32 |
| | 300 | 0 | 0.00 | 0.00 | 0.33 | 0.33 | 0.00 | 0.00 | 0.24 | 0.24 |
| | 0 | 1 | 0.62 | 0.63 | 3.22 | 3.22 | 0.43 | 0.42 | 0.57 | 0.57 |
| | 50 | 1 | 0.49 | 0.49 | 2.88 | 2.90 | 0.22 | 0.21 | 0.54 | 0.54 |
| | 150 | 1 | 0.10 | 0.10 | 2.00 | 1.98 | 0.02 | 0.02 | 0.45 | 0.44 |
| | 300 | 1 | 0.01 | 0.01 | 0.62 | 0.61 | 0.01 | 0.01 | 0.28 | 0.28 |
| | 0 | 3 | 0.54 | 0.53 | 3.23 | 3.21 | 0.36 | 0.36 | 0.58 | 0.58 |
| | 50 | 3 | 0.58 | 0.57 | 3.37 | 3.35 | 0.48 | 0.48 | 0.58 | 0.58 |
| | 150 | 3 | 0.29 | 0.28 | 2.53 | 2.51 | 0.05 | 0.05 | 0.49 | 0.50 |
| | 300 | 3 | 0.02 | 0.02 | 1.16 | 1.15 | 0.01 | 0.01 | 0.35 | 0.35 |
